# Mouse behaviour on the trial-unique non-matching-to-location (TUNL) touchscreen task reflects a mixture of distinct working memory codes and response biases

**DOI:** 10.1101/2022.10.30.514444

**Authors:** Daniel Bennett, Jay Nakamura, Chitra Vinnakota, Elysia Sokolenko, Jess Nithianantharajah, Maarten van den Buuse, Nigel C. Jones, Suresh Sundram, Rachel Hill

**Affiliations:** School of Psychological Sciences, Monash University, Melbourne, VIC 3180, Australia; Department of Psychiatry, Monash University, Melbourne, VIC 3180, Australia; Laboratory for Molecular Mechanisms of Brain Development, RIKEN Center for Brain Science, Saitama, Japan; Discipline of Anatomy and Pathology, School of Biomedicine, University of Adelaide, Adelaide SA 5005, Australia; The Florey Institute of Neuroscience and Mental Health, Melbourne, VIC 3052, Australia; School of Psychology and Public Health, La Trobe University, Melbourne, VIC 3086, Australia; Department of Neuroscience, Central Clinical School, Monash University, Melbourne, VIC 3004, Australia; Department of Neurology, The Alfred Hospital, Commercial Road, Melbourne, VIC 3004, Australia; Department of Medicine, Royal Melbourne Hospital, The University of Melbourne, Melbourne, VIC 3052, Australia; Mental Health Program, Monash Health, Clayton, VIC 3168, Australia

## Abstract

The trial-unique non-matching to location (TUNL) touchscreen task shows promise as a translational assay of working memory deficits in disorders including autism, ADHD, and schizophrenia. Although it is commonly assumed that the TUNL task predominantly measures spatial working memory in rodents, this proposition has not previously been tested. In this project, we used computational modelling of behaviour from mice performing the TUNL task (total *N* = 163 mice across three datasets; 158,843 total trials). Contrary to common assumptions, behaviour on the TUNL task did not exclusively reflect spatial working memory. Instead, choice behaviour was explained as a mixture of both retrospective (spatial) working memory and prospective working memory for an intended behavioural response, as well as animal-specific response biases. We suggest that these findings can be understood within a resource-rational cognitive framework, and propose several task-design principles that we predict will maximise spatial working memory and minimise alternative behavioural strategies.

## Introduction

Working memory (WM)—the active maintenance and manipulation of information within a transient, limited-capacity memory store—is crucial for many of the tasks that humans face in their day-to-day lives (Baddeley, 2010; Cowan, 2014; Oberauer & Lin, 2017). For example, WM allows a person to maintain a restaurant’s address in mind as they drive across town to dinner (Blacker et al., 2017), and also underpins the mental arithmetic that they perform after the meal as they split the bill with friends (DeStefano & LeFevre, 2004). WM is impaired across a range of psychiatric disorders, including schizophrenia, schizoaffective disorder, bipolar disorder and psychosis (Gold et al., 2019), attention-deficit/hyperactivity disorder (Kofler et al., 2018), and autism (Steele et al., 2007; Zhang et al., 2020). Moreover, WM deficits are predictive of poor clinical and functional outcomes in both schizophrenia (Fu et al., 2017; Jenkins et al., 2018) and autism (Leung et al., 2016; Troyb et al., 2014).

Despite this, there are no current treatments that address WM impairments in psychiatric disorders. In overcoming this barrier, animal models are key to trialling novel therapeutic compounds (Castner et al., 2004; Dudchenko et al., 2013). However, as is the case with many disorders of the brain, the translation of novel therapeutics to clinical application is dependent on the validity of the animal model used in pre-clinical evaluations of potential therapeutics (Pound & Ritskes-Hoitinga, 2018).

Behaviours indicative of human-like WM have been documented across a wide range of species, including insects, fish, birds, rodents, and non-human primates (Aultman & Moghaddam, 2001; Bloch et al., 2019; Giurfa et al., 2001; Miller et al., 1996; Roberts, 1972). However, this similarity does not necessarily entail that WM operates in the same way across species. In fact, there are pronounced between-species differences in WM, particularly in the apparent capacity of the WM store and the rate at which information is forgotten during a retention interval (Carruthers, 2013; Lind et al., 2015; Roberts & Santi, 2017). From a translational perspective, this raises a critical question: are the neurocognitive processes that underlie WM in a given species sufficiently similar to the processes underlying human WM to use that species as a model system?

### The TUNL task

Recent research initiatives have sought to develop cognitive tasks for use in animal models that maximise the similarity between animal and human WM tasks (e.g., as part of the CNTRICS initiative; Carter et al., 2011). The particular focus of this research has been on rodent WM, with the goal of producing more reliable models for preclinical drug testing (Hvoslef-Eide et al., 2015). A particularly promising task is the trial-unique non-matching-to-location (TUNL) task, a touchscreen-based task that allows for high-throughput measurement of spatial WM in mice and rats (Bussey et al., 2012; Kim et al., 2015; Oomen et al., 2013; Talpos et al., 2010). In the TUNL task, animals are trained to remember the location of a ‘sample’ stimulus (an illuminated square on the touchscreen) during a retention interval in which the sample location is no longer illuminated. In a choice phase, animals are shown two illuminated squares and are rewarded if they touch the square that does not match the sample stimulus (see Figure 1A).

**Figure 1.**
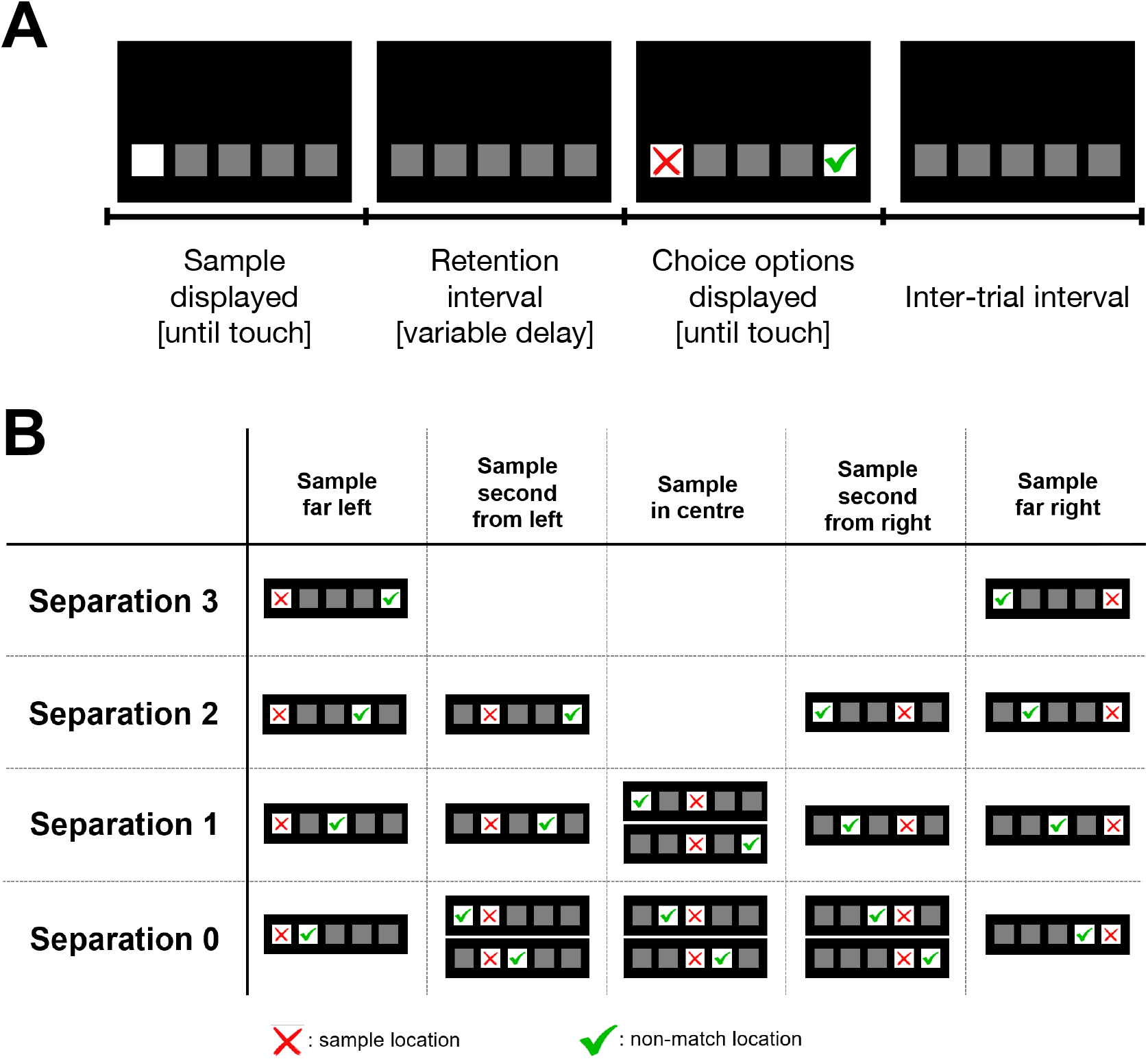
Trial schematic for the mouse TUNL task. **A:** Sequence of events in each trial. The animal initiates the delay period by touching the sample location (white square). After the retention interval, the animal is shown two lit squares; to receive a reward it must touch the square that is a non-match for the trial’s sample location (correct location: green tick; incorrect location: red cross; ticks and crosses are for illustration purposes only and are not visible to mice). **B**. Overview of each of the 20 possible configurations of sample and non-match location. Separation conditions are defined in terms of the number of unlit squares that separate the sample location and the non-matching location (e.g., in a Separation 3 trial, the sample location and the non-match location are separated by three unlit squares). Within each separation type, there are multiple different configurations of sample locations (red crosses) and non-match locations (green ticks). The 20 different possible configurations of sample location and non-match location that can be presented comprise two configurations constituting a Separation 3 trial, four for Separation 2 trials, six for Separation 1 trials, and 8 for Separation 0 trials.

The TUNL task uses a non-match-to-sample paradigm rather than a match-to-sample paradigm (in which the matching stimulus is selected after a delay; e.g., Blough, 1959) because animals performing match-to-sample tasks have frequently been found to use compensatory behavioural strategies rather than solving the task by means of WM. Specifically, match-to-sample paradigms allow the animal to predict at the start of the retention interval what response it should make in future. In these circumstances, animals frequently adopt ‘overt rehearsal’ strategies, such as orienting towards the correct response option or repeatedly operating a correct response lever during the retention interval (see, e.g., Dudchenko & Sarter, 1992; Hearst, 1962; Mackintosh, 1983). This confound is thought to be abolished in the TUNL by removing any incentive for the animal to adopt a compensatory behavioural strategy and theoretically making the task a purer measure of WM.

However, a separate and more subtle question, which has not been considered with respect to the TUNL task to date, concerns the nature of the representational code with which information is held in WM. Broadly, we can distinguish two kinds of representational codes for WM: a *retrospective code*, in which animals maintain a representation of a previous stimulus, and a *prospective code*, in which animals maintain a representation of an anticipated future behavioural response (cf. Cook et al., 1985; Roitblat, 1982). In the TUNL task, retrospective coding would involve encoding the spatial location of the sample stimulus and retaining this location in WM during the retention interval. By contrast, prospective coding would involve observing the spatial location of the sample and pre-emptively encoding a planned behavioural response: for example, if the sample stimulus were the left-most box, then prospective coding would entail the animal maintaining in their WM an intention to choose whichever response option is further to the right in the choice phase.

To our knowledge, all work on the TUNL to date has assumed that animals adopt a retrospective WM code in this task (i.e., that they encode and maintain in WM the *spatial location* of the sample stimulus). This is a reasonable hypothesis *a priori*, since a retrospective WM code maximises reward on the task, and is thought to be how humans solve similar WM tasks (e.g., Della Sala et al., 1999). However, although humans are thought to use a primarily retrospective memory code in standard visuospatial WM tasks (e.g., Della Sala et al., 1999; Glahn et al., 2003; McCarthy et al., 1994), non-human animals have been shown to use a mixture of prospective and retrospective memory codes in other, related memory tasks (Cook et al., 1985; Ferbinteanu & Shapiro, 2003; Kametani & Kesner, 1989; Kesner, 1989; Zimmer, 2008). It has not yet been determined what WM coding and behavioural strategies are employed by animals on the TUNL task (but see Voitov & Mrsic-Flogel, 2022). This is particularly pertinent given that it is possible for animals to achieve above-chance performance on the TUNL task solely using prospective WM^1^.

The touchscreen behavioural tasks generate large amounts of trial-unique data that is overlooked when only analysing summary measures (e.g., response accuracy to all trial types). In this project, we sought to take advantage of the rich data provided by touchscreen behavioural tasks to investigate the behavioural strategies and WM coding of mice performing the TUNL task. The primary tool that we used to answer this question was computational modelling of behaviour, which provides a powerful method for deconstructing animal behaviour into its constituent cognitive subcomponents (see, e.g., Broschard et al., 2019, 2021; Doll et al., 2012; Langdon et al., 2019). We compared a suite of computational models that quantified the degree to which mice’s behaviour reflected two distinct WM codes: retrospective WM (i.e., memory for spatial location of sample stimulus) vs. prospective WM (i.e., WM for intended response direction), as well as two idiosyncratic response biases: side biases (i.e., preferring to respond at the left/right, independent of choice options) and distal-response bias (i.e., preferring to respond at locations closer to the edges of the testing arena). These choice biases are animal-specific preferences for different response locations that are independent of the location of the sample stimulus and static across time. Our results demonstrate that, contrary to common assumptions, mouse behaviour on the TUNL task does not solely reflect retrospectively-coded (i.e., spatial) WM. Instead, we found evidence for a mixture of both kinds of WM coding systems, as well as for both kinds of choice biases.

## Results

### Overview of task and data

We analysed behavioural data from mice performing the TUNL task in touchscreen operant chambers (Figure 1; see Materials and Methods for a full task description). Here we report data from a total sample of 158,843 choice trials from *N* = 163 mice across three distinct datasets (see Table 1), focusing our analyses on the post-training ‘probe’ test phase of the task, which is typically taken as the measure of spatial WM performance in studies using the TUNL task (e.g., Kim et al., 2015; Nilsson et al., 2016; Zeleznikow-Johnston et al., 2017). Note that, although the three datasets had similar training protocols, each of the three datasets used a slightly different set of trial types in the probe phase of the task. We deliberately selected datasets with a heterogeneous set of trial types because our overall goal was to ensure that our model selection procedures were broadly generalisable. By fitting models to datasets that differed in their trial types and probe-phase parameters, we can be confident that the overall best-fitting model was not simply one that was over-fit to a specific set of trial types or testing parameters.

**Table 1.** Overview of datasets

| Dataset # | Original publication | $N$ mice (female /male) | $N$ probe-phase trials | Separation conditions | Delays (seconds) | Correction trials at test? |
| --- | --- | --- | --- | --- | --- | --- |
| 1 | Nakamura et al. (2021) | 83 (46/37) | 113,599 | S0, S1, S2, S3 | 0, 3, 6, 9, 12, 15, 18, 21, 24 | No |
| 2 | Vinnakota et al. (in prep.) | 44 (19/25) | 24,947 | S1, S2, S3 | 1, 2, 3 | Yes |
| 3 | Sokolenko et al. (2020) | 36 (0/36) | 20,297 | S1* | 1, 2, 3, 6 | Yes |

\* Dataset 3 only tested S1 trials for which either the sample location or the non-match location was at the centre response location (Figure 1B: Separation-1 row, columns 1, 3, and 5).

Each separation condition in the TUNL task can be constructed using multiple different configurations of sample location and non-match location: a Separation-3 (or S3) trial, for instance, can be constructed either with the sample at the far left and the (correct) non-match option at the far right, or vice versa. This is a strength of the task because, unlike earlier non-match-to-sample tasks involving only two response levers (e.g., Chudasama & Muir, 1997), there are 20 different unique configurations of sample and non-match location that can be tested across the four different separation conditions (see Figure 1B). This relative increase in trial-uniqueness is thought to reduce the likelihood of mediating behavioural responses (e.g., repeatedly pressing a response lever during the retention interval; Talpos et al., 2010).

### Different configurations at the same spatial separation are not interchangeable

In previous studies, behaviour on the TUNL task has been analysed by averaging across the different possible stimulus configurations within a separation condition to create a single summary measurement per animal per separation and delay (i.e., averaging across the different configurations within each row of Figure 1B). This summary-statistic approach implicitly assumes that different stimulus configurations are otherwise interchangeable with one another, and that the only feature of a sample/non-match location pair that is relevant for behaviour is the spatial separation between the two locations. We first tested whether this assumption was met in our data by testing for differences in performance across different configurations within a separation condition.

As can be seen in Figure 2, this assumption was not met: instead of different configurations falling on the same temporal decay curve, as we would expect if configurations were interchangeable with one another, there was marked heterogeneity between configurations within the same separation condition. Mixed-effects logistic regression analyses revealed that these differences were statistically significant in all three of our datasets: for Dataset 1 (Figure 2A), there was a significant difference between different configurations on accuracy for S0 trials (χ^2^(7) = 287.26, *p* < .001), S1 trials (χ^2^(5) = 402.39, *p* < .001), and S2 trials (χ^2^(3) = 170.14, *p* < .001), though the effect of configuration was not statistically significant for S3 trials (χ^2^(1) = 1.67, *p* = .20). For Dataset 2 (Figure 2B), there was a significant effect of configuration on accuracy for S1 trials (χ^2^(5) = 21.18, *p* < .001) and S2 trials (χ^2^(3) = 23.06, *p* < .001), though again the effect of configuration was not statistically significant for S3 trials (χ^2^(1) = 0.08, *p* = .77). For Dataset 3 (Figure 2C), which used a smaller number of unique trial configurations, the effect of configuration on accuracy for S1 trials (χ^2^(3) = 5.79, *p* = .12) was not statistically significant, but there was a significant interaction between configuration and delay (χ^2^(3) = 19.96, *p* < .001), indicating that the rate at which performance deteriorated as a function of delay differed between the different configurations. These results indicate that, contrary to common assumptions, there are widespread differences in TUNL performance depending on the exact configuration of sample location and non-match location that was tested.

**Figure 2.**
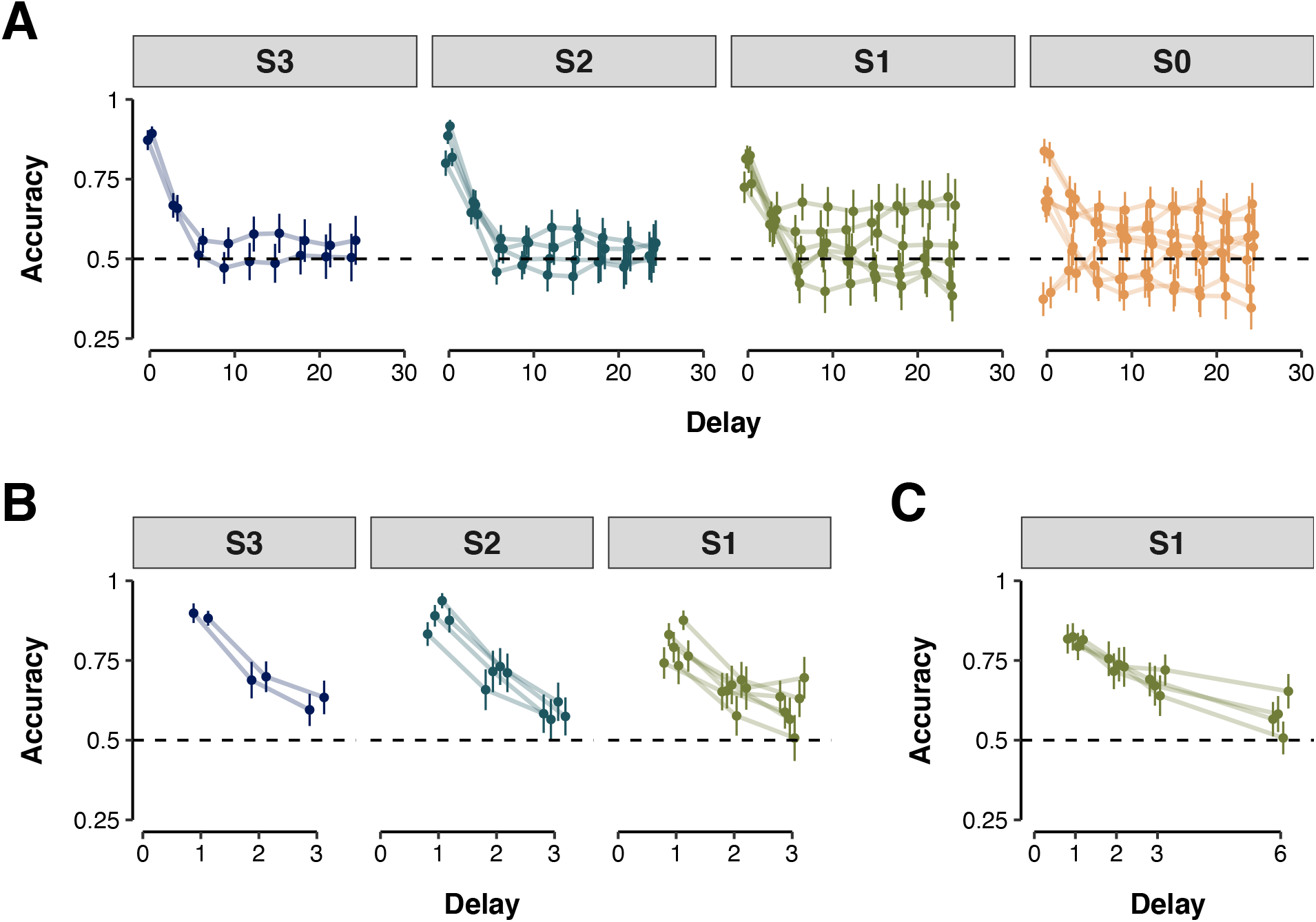
Mouse behavioural data as a function of separation and delay across three independent datasets (**A**: Nakamura et al., 2021; **B**: Vinnakota et al., in prep.; **C**: Sokolenko et al., 2020). Each subplot depicts mean proportion of correct responses (y-axis) across animals as a function of delay between memory sample and response screen (x-axis) and separation condition (plot facets). Each individual point within a delay/separation condition presents data from a unique configuration of sample location and non-match location (see Figure 1B). Error bars represent the 95% confidence interval of the mean.

### Between-configuration differences are consistent with prospective WM coding

We next sought to unravel the source of these between-configuration differences in behaviour. In particular, we investigated a striking pattern in which some pairs of Separation-0 stimulus configurations showed markedly different patterns of performance from one another even though they shared the same pair of response options in the choice phase of the trial.

One such pair (from Dataset 1) is presented in Figure 3A. When the sample location was at the far left and the correct non-match response location was second from the left, animals performed the task well above chance-level performance at zero delay (mixed-effects logistic regression; intercept = 1.26, *p* < .001), and this performance level slowly deteriorated with increases in the duration of the retention interval (*β* = -0.05, χ^2^(1) = 22.99, *p* < .001). Notably, however, when these locations were reversed (sample location second from left and correct non-match response location at far left) animals performed the task significantly *below* chance performance at zero delay (intercept = -0.49, *p* < .001), and performance *improved* with increases in the duration of the retention interval (*β* = 0.02, χ^2^(1) = 5.99, *p* = .01). This pattern was not restricted to the left side of the testing arena: a similar pattern was observed for corresponding Separation-0 configurations on the right of the testing arena (see Supplementary Figure S2).

**Figure 3.**
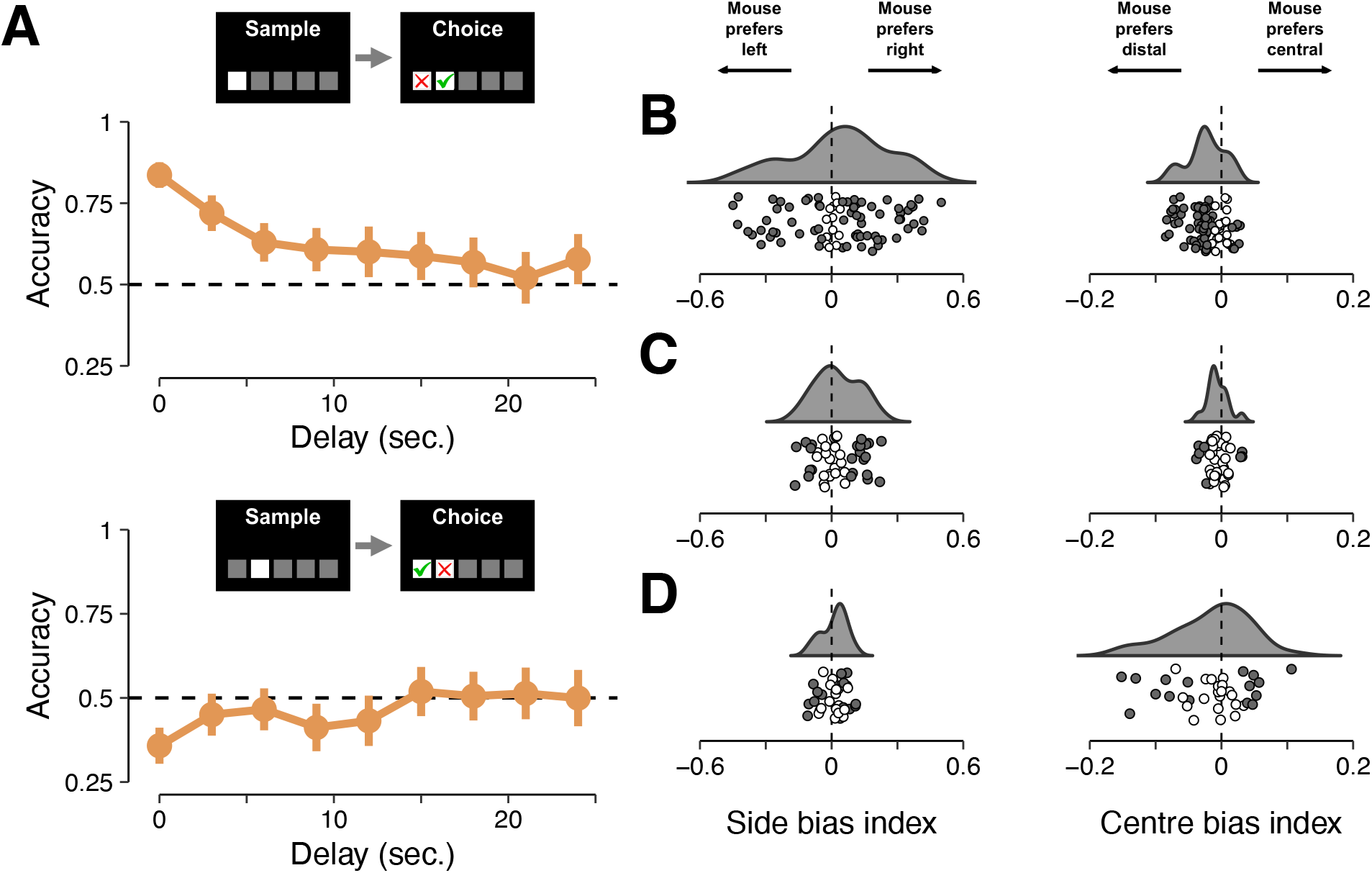
**(A)** Demonstration of direction-dependent effects in two putatively similar S0 choice configurations across multiple delays from Dataset 1. Top: sample location [red cross] at far left and non-match location [green tick] second from the left; mice display above-chance performance at delay 0 that deteriorates with increasing delay duration. Bottom: sample location second from the left and non-match location at the far left; mice show below-chance performance at delay 0, and improving performance as delays increase. Error bars represent the 95% confidence interval of the mean. **(B-D)** Side-biases (left column) and centre-biases (right column) for Dataset 1 **(B)**, Dataset 2 **(C)**, and Dataset 3 **(D)**. A positive side bias index indicates that a mouse preferred to respond at the right hand side of the chamber, and a negative side bias index indicates that a mouse preferred to respond at the left hand side of the chamber. A positive centre bias indicates that a mouse preferred to respond at the centre of the chamber, and a negative centre bias indicates that a mouse preferred to respond at the edges of the chamber. Distributions represent estimates of the group-level distributions in each dataset. Individual points depict statistics for individual animals. Filled datapoints represent animals with a biases that were significantly different from chance; unfilled datapoints represent animals with biases not significantly different from chance.

This striking pattern of results is difficult to reconcile with the standard hypothesis that animals are performing this task using spatial WM (i.e., a retrospective memory code; maintaining a representation in WM of the spatial location of the sample stimulus). Under this standard explanation, the difficulty of Separation-0 trials results from the strong spatial similarity in the two possible response locations in the choice phase locations. Crucially, however, this similarity is symmetric: by definition, the spatial similarity of the far-left location compared to the second-from-left location is identical to the spatial similarity of the second-from-left location compared to the far-left location. Because of this symmetry, it is difficult to explain under the assumption of retrospective WM coding how behaviour could be significantly above-chance (and deteriorating with increasing delays) in the top panel of Figure 3A but significantly below-chance (and improving with increasing delays) in the bottom panel of Figure 3A.

By contrast, this puzzling pattern of effects is relatively easily explained if we instead assume that animals were using prospective WM coding. Recall that under a prospective WM code, animals would maintain in WM during the retention interval a behavioural intention. This is possible because—depending on the sample location—the animal can often predict with reasonable accuracy the direction in which it should respond on the basis of the sample location that it sees. For instance, if the sample location is at the far left, then the animal should always choose whichever response option is further to the right at the choice phase, independent of separation condition (left column of Figure 1B and Supplementary Figure S1). Similarly, if the sample location is second from the left, then in 75% of trials the correct response location will be whichever choice option is further to the right in the choice phase (second column from left in Figure 1B and Supplementary Figure S1); this means that the TUNL task can be partially solved using prospective WM coding. Then, if the animal is using a prospective WM code, when it observes a sample location second from the left (Figure 3A, bottom panel) it will encode in WM a prospective intention to choose whichever response option is further right, which will produce below-chance performance in this particular case. Moreover, its performance will also improve with increasing delays for this configuration because the animal’s choices will become more random (and therefore more correct in this case, until it reaches chance level) as the prospective WM representation fades.

### Animals also show idiosyncratic side biases and distal-response biases

We also investigated how behaviour in the TUNL task was influenced by two response biases unrelated to WM: side biases and distal-response biases. Side biases refer to the tendency for individual animals to prefer leftward or rightward response options, and have been observed across species and operant conditioning paradigms (Alber & Strupp, 1996; Miletto Petrazzini et al., 2020). By contrast, distal-response biases^2^ are more specific to the TUNL task, and refer to the tendency of mice to prefer response locations closer to the left or right walls of the testing arena over more central response locations (see Kim et al., 2015).

We found evidence for both side biases and distal-response biases in all three datasets (Figure 3B-D). For side biases (Figure 3, centre column), although there was no average preference for leftward versus rightward responses at the group level in any individual dataset (Dataset 1: *t*(82) = 1.63, *p* = .11; Dataset 2: *t*(43) = 1.80, *p* = .08; Dataset 3: *t*(89) = 1.23, *p* = .23), permutation tests revealed statistically significant side biases within the behaviour of a majority of individual animals in each dataset. In Dataset 1, 27/83 mice (33%) showed a significant leftward bias and 43/83 mice (52%) showed a significant rightward bias; in Dataset 2, 8/44 mice (18%) showed a significant leftward bias and 15/44 mice (34%) showed a significant rightward bias; in Dataset 3, 7/36 mice (19%) showed a significant leftward bias and 12/36 mice (33%) showed a significant rightward bias. Across all datasets, there was no evidence for an association between the strength of animals’ side biases and their accuracy rate on the task (Dataset 1: Pearson *r*(83) = -.08, *p* = .49; Dataset 2: *r*(44) = -.17, *p* = .27; Dataset 3: *r*(36) = -.12, *p* = .48).

Distal-response biases (Figure 3, right column) were less prominent overall than side biases, but nevertheless accounted for a proportion of group-level and animal-level variance. At a group-level, we found a significant preference for distal response options over central response options in both Dataset 1 and Dataset 2 (Dataset 1: *t*(82) = -7.27, *p* < .001; Dataset 2: *t*(43) = -2.55, *p* = .01), though not in Dataset 3 (Dataset 3: *t*(35) = -1.56, *p* = .12). Moreover, permutation tests revealed statistically significant distal- or central-response biases within the behaviour of a sizeable proportion of individual animals in each dataset. In Dataset 1, 52/83 mice (63%) showed a significant distal-response bias and 9/83 mice (11%) showed a significant centre-response bias; in Dataset 2, 5/44 mice (11%) showed a significant distal-response bias and 3/44 mice (7%) showed a significant centre-response bias; in Dataset 3, 8/36 mice (22%) showed a significant distal-response bias and 7/36 mice (19%) showed a significant centre-response bias. It is noteworthy that, even though there was a significant group-level preference for distal response options on average, some individual animals showed a significant preference for the centre response options. This speaks to the heterogeneity of behaviour in the TUNL task and raises the possibility that individual animals might perform the task using different combinations of WM coding schemes and response biases. In Datasets 1 and 2, there was no evidence for an association between the strength of animals’ distal-response biases and their accuracy rate on the task (Dataset 1: Pearson *r*(83) = .03, *p* = .82; Dataset 2: *r*(44) = .01 *p* > .99). In Dataset 3, animals that preferred central response options more strongly tended to perform better on the task overall (*r*(36) =.44, *p* < .01). Given that Dataset 3 tested only S1c trials (i.e., trials in which either the sample location or the non-matching location was in the centre), this suggests that animals that were better able to suppress the instinct to move towards the walls of the testing chamber when the correct response location was in the centre might have performed better on the task overall.

### Task behaviour is best explained by a mixture of WM codes and response biases

The results above provide evidence that, in addition to any effects of retrospective WM, choice behaviour on the TUNL task was also affected by prospective WM, side biases, and distal-response biases. To explicate the relative importance of each of these different *response factors*, we next turned to computational modelling of animals’ choice behaviour.

A full description of all computational models and the hierarchical Bayesian framework within which they were fit to data is available in the Materials and Methods section. Briefly, all models assumed that choices on a given trial were driven by the competition between two response strengths: one for the matching location and one for the non-matching location. We compared a set of different computational models that differed in their assumptions about how the response strengths for each response location was determined. The specific set of models that we compared comprised all one-, three-, and four-way combinations of the four response factors (see Table 2 and Materials and Method).

**Table 2.** Overview of computational model fits to each dataset

| Model number | N model parameters per mouse | Forgetting function | Retrospective working memory | Prospective working memory | Side biases | Distal-response biases | Dataset 1 |  | Dataset 2 |  | Dataset 3 |  |
| --- | --- | --- | --- | --- | --- | --- | --- | --- | --- | --- | --- | --- |
| | | | | | | | WAIC | ΔWAIC (SE) | WAIC | ΔWAIC (SE) | WAIC | ΔWAIC (SE) |
| M1 | 3 * | Power-law | ✓ | - | - | - | 71,495.2 | 15,678.6 (362.0) | 7,940.5 | 584.8 (74.3) | 3807.6 | 441.9 (76.1) |
| M2 | 2 | Power-law | - | ✓ | - | - | 71,471.2 | 15,654.7 (361.1) | 8,491.5 | 1,135.8 (103.5) | 6753.4 | 3,387.6 (359.0) |
| M3 | 1 | N/A | - | - | ✓ | - | 66,937.5 | 11,120.9 (370.9) | 14,189.0 | 6,833.3 (300.2) | 10,405.6 | 7,039.8 (576.7) |
| M4 | 1 | N/A | - | - | - | ✓ | 76,250.8 | 20,434.3 (422.7) | 14,321.4 | 6,965.7 (296.0) | 10,502.6 | 7,136.8 (571.0) |
| M5 | 4 | Power-law | - | ✓ | ✓ | ✓ | 56,025.8 | 211.3 (65.5) | 7,378.2 | 22.5 (15.7) | 3,812.1 | 446.3 (80.9) |
| M6 | 5 * | Power-law | ✓ | - | ✓ | ✓ | 57,088.1 | 1,271.6 (117.6) | 7,511.2 | 155.5 (36.7) | 3,382.2 | 16.5 (18.5) |
| M7 | 5 * | Power-law | ✓ | ✓ | - | ✓ | 68,261.1 | 12,444.5 (335.2) | 7,771.4 | 415.7 (61.4) | 3,707.8 | 342.0 (69.7) |
| M8 | 5 * | Power-law | ✓ | ✓ | ✓ | - | 58,258.4 | 2,441.9 (148.0) | 7,511.9 | 156.2 (36.7) | 3,401.1 | 35.3 (17.4) |
| M9 | 6 * | Power-law | ✓ | ✓ | ✓ | ✓ | **55,816.5** | - | **7,355.0** | - | **3,365.8** | - |

Note: WAIC values are presented on a deviance scale (lower values indicate better model fit).
\* One fewer parameter per mouse in Dataset 3 because a smaller number of separations in this dataset rendered the $\alpha$ parameter non-identifiable (see Method)

Overall, in all three datasets the best-fitting model was model M9, in which behaviour was produced by a mixture of all four response factors (see Supplementary Table S3 for parameter estimates). This indicates that in each of the datasets that we considered, the model that best explained the data was one in which both prospective and retrospective WM influenced animals’ choices, and in which individual animals held both side biases and distal-response biases. The one caveat to this otherwise consistent result comes from Dataset 3: although model M9 was still the best-fitting model overall as measured by the WAIC statistic, model M6 provided a statistically equivalent fit to data when model fit uncertainty was accounted for (i.e., the difference in WAIC values between M6 and M9 in Dataset 3 was smaller than the standard error of this difference; cf. Bennett et al., 2021; Weber et al., 2022).

As well as being the best-fitting model in relative terms, model M9 provided an excellent absolute fit to data in all three datasets (see posterior predictive checks in Figure 4). This implies that the four response factors in this model are each necessary to explain some portion of behavioural variance. However, this does not imply that all four factors accounted for an equal amount of variance in behaviour. To investigate this more quantitatively, we computed the amount of unique variance explained by each of the four response factors in M9 (see Table 3). The results of this analysis indicate substantial heterogeneity across datasets in the proportion of variance accounted for by different factors. In particular, the balance between retrospective and prospective WM differed substantially across datasets: in Dataset 1, prospective WM uniquely accounted for 17.9% of group-level behavioural variance and retrospective WM only 3.1%, whereas in Dataset 3 prospective WM uniquely accounted for only 6.7% of variance whereas retrospective WM accounted for 22%.

**Figure 4.**
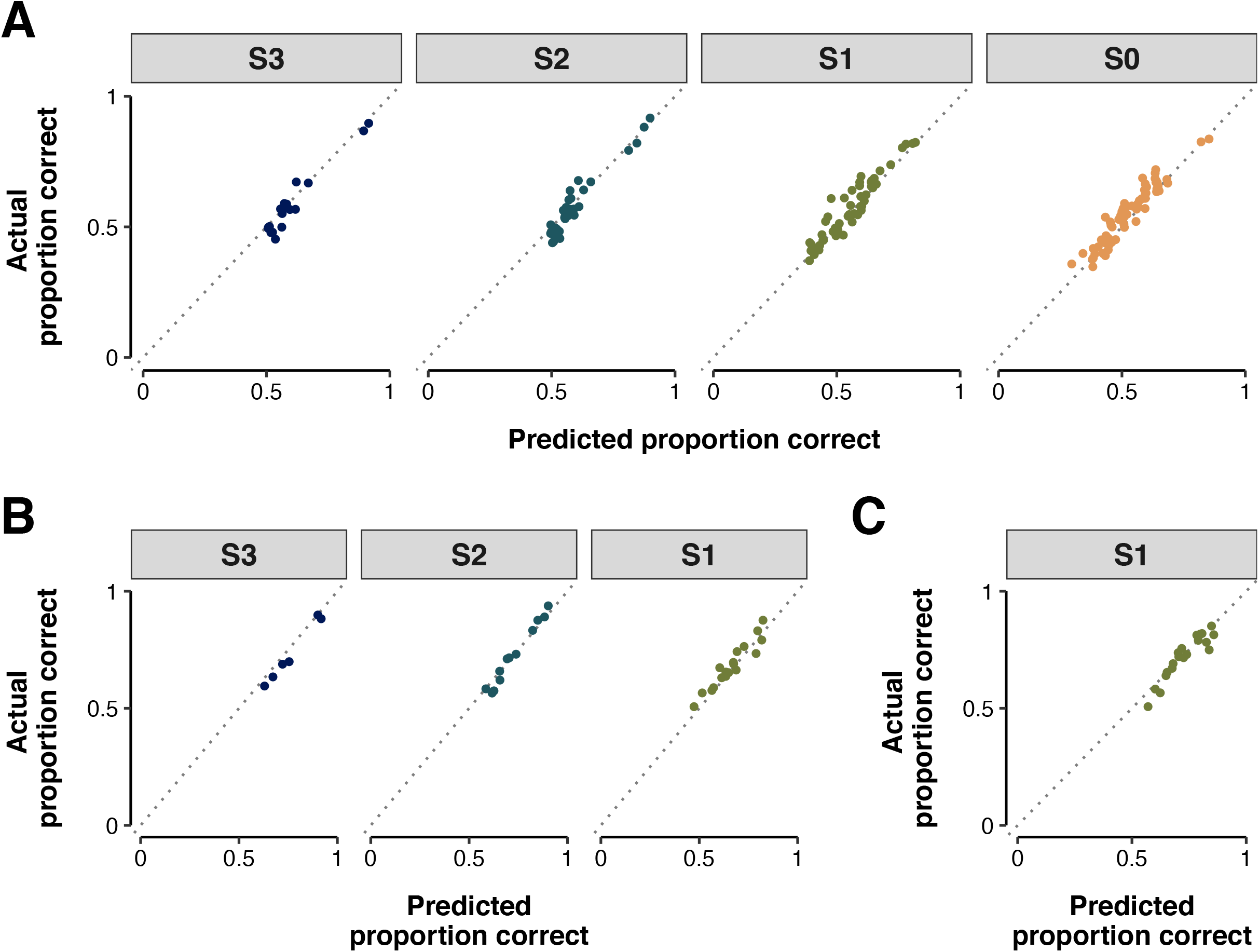
Posterior predictive checks for the best-fitting computational model on each of the three datasets (**A:** Dataset 1; **B:** Dataset 2; **C**: Dataset 3). Subplots show the correspondence between the actual mean proportion of correct responses across mice (y-axis) and the model’s predicted mean proportion of correct responses across mice (x-axis), with different separation conditions presented in different plot facets and colours. Each point represents a unique combination of sample location, non-match location, and delay, averaged across mice. The diagonal dotted line represents equality between actual and predicted proportion; in a perfectly calibrated model, every point would fall exactly on this diagonal. The best-fitting computational model provides an excellent fit to data in each of the three independent datasets.

**Table 3.** Partitioning of unique variance explained within model M9

| Dataset # | Overall<br>model R<sup>2</sup> | Unique variance explained by each response factor * |  |  |  |
| --- | --- | --- | --- | --- | --- |
|  |  | Spatial/<br>retrospective<br>working memory | Prospective<br>working memory | Distal-response<br>biases | Side<br>biases |
| 1 | 88.6% | 3.1% | 17.9% | 32.8% | 4.6% |
| 2 | 91.1% | 3.9% | 4.5% | 7.5% | 2.8% |
| 3 | 85.6% | 22.0% | 6.7% | 7.6% | 1.7% |

\* Calculated as change in group-level R<sup>2</sup> when removing a response factor from the model.

### Individual animals varied in their mixture of WM codes and response biases

The results presented above provide evidence that, at a group level, animal behaviour was best explained as a mixture of WM codes and response biases. Of particular note, there was evidence that mouse behaviour in each of the three datasets was produced by *both* prospective and retrospective WM. There are several different ways that this pattern of data might emerge: first, each individual mouse might have used *either* prospective *or* retrospective WM, and each dataset might have consisted of mixtures of these two kinds of mouse in varying proportions. Second, such a result might also arise if individual animals used both prospective and retrospective WM coding. We next sought to determine which of these two possibilities provided the best account of data for individual animals (see Figure 5).

**Figure 5.**
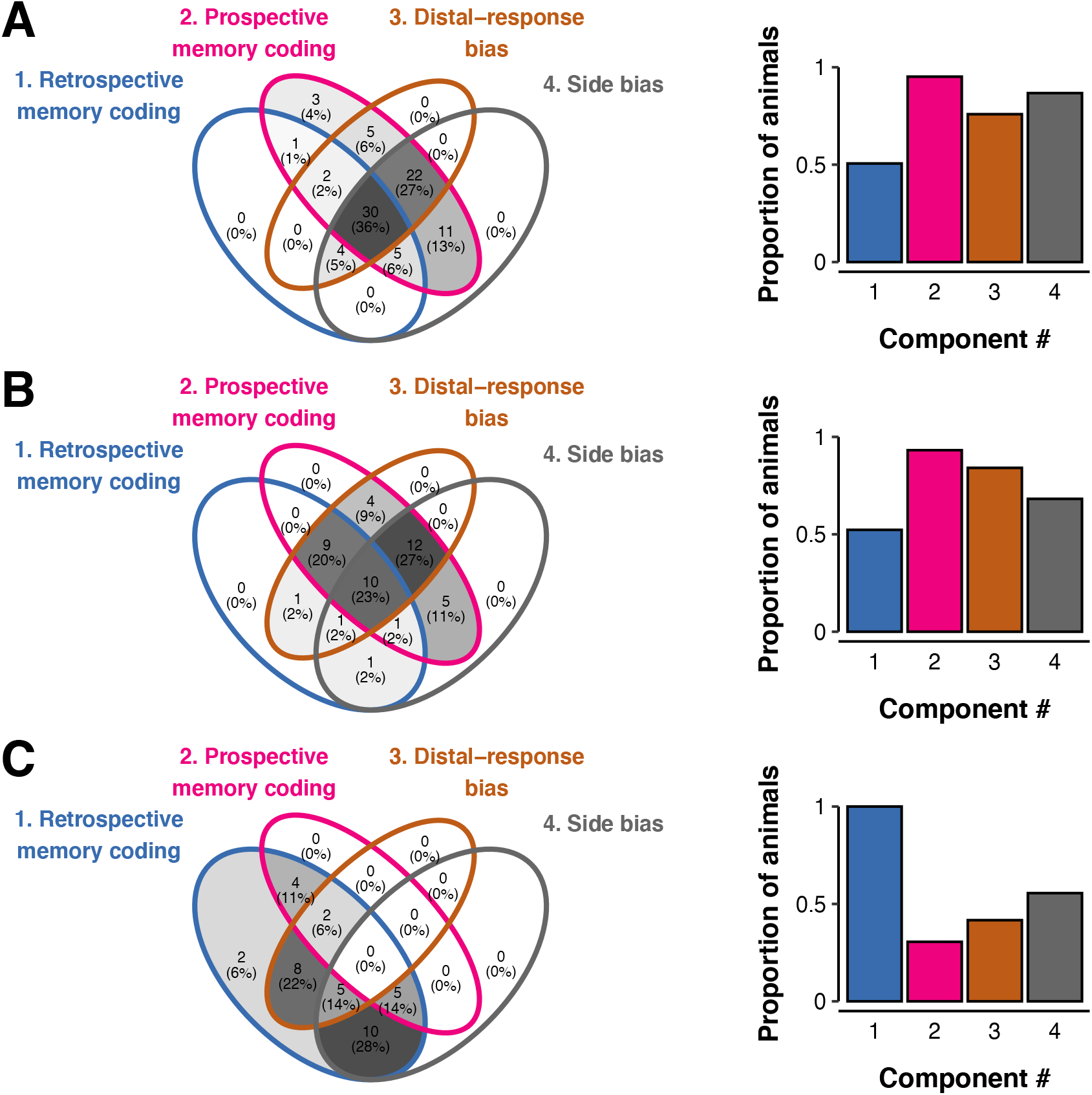
Breakdown of significant model components across mice in each dataset (**A:** Dataset 1; **B**: Dataset 2; **C**: Dataset 3). Venn diagrams present the breakdown of the numbers of mice who fit into each possible configuration of significant model components (presented as raw numbers and percentage of dataset). Histograms depict the proportion of mice in each dataset who displayed a significant effect of each model component overall (i.e., the sum of the percentages within each coloured circle in the Venn diagrams).

**Figure 5.**
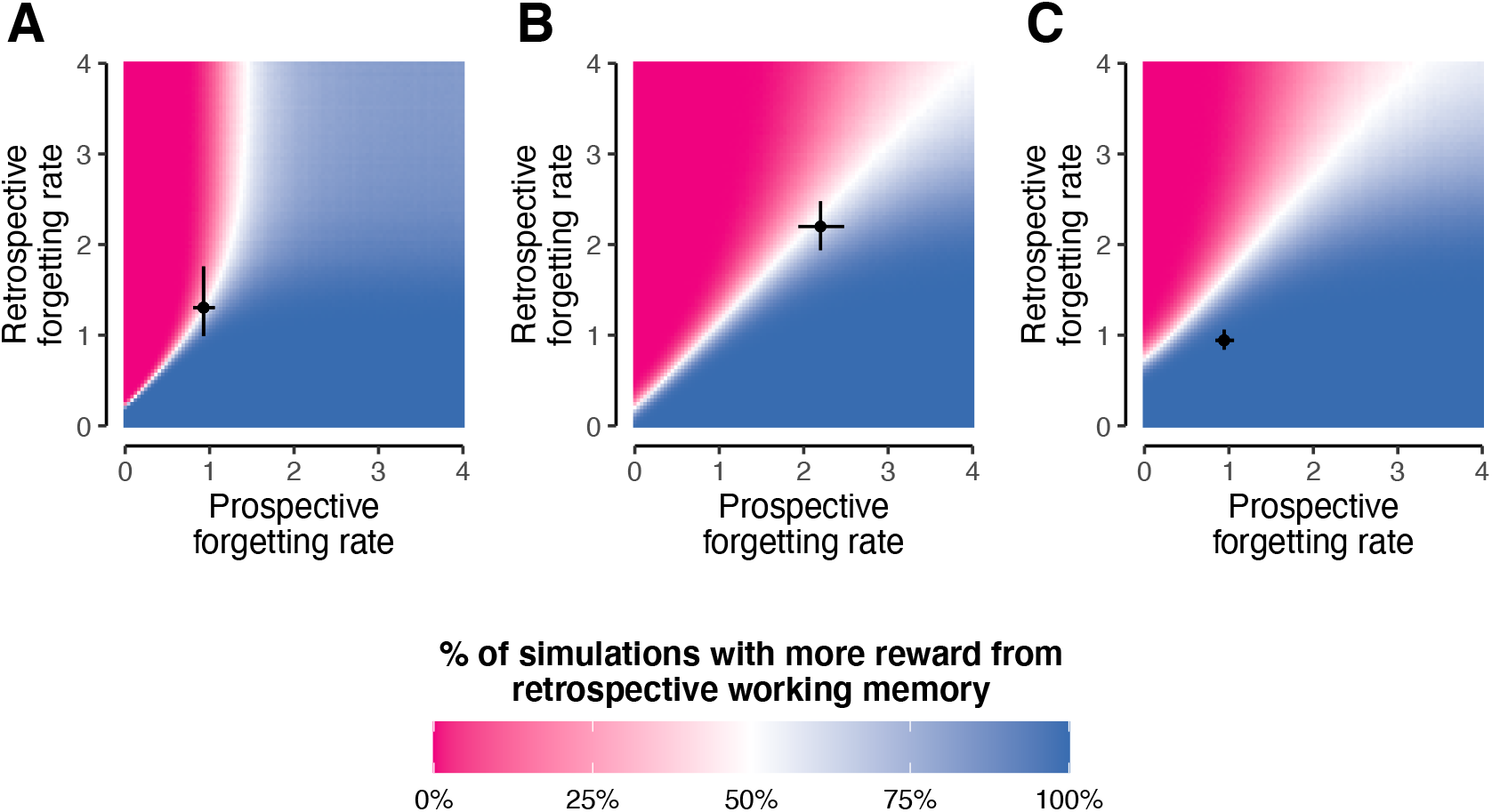
Simulated performance under different forgetting rates for prospective and retrospective WM in the best fitting model for each dataset (**A**: trial types and delays as per Dataset 1; **B**: trial types and delays as per Dataset 2; **C**: trial types and delays as per Dataset 3). Blue (/pink) regions of each heatmap denote parameter regimes under which more reward can be acquired using retrospective (/prospective) WM. White regions denote parameter regimes in which retrospective and prospective WM produce equivalent performance on average. Superior performance for prospective WM typically occurs when the forgetting rate for prospective WM is markedly lower than the forgetting rate for retrospective WM, though comparison of the three panels indicates that the exact region of superiority depends on the trial types that are tested. Points and error bars in each dataset denote the mean estimated forgetting rate for each WM type in each dataset +/-the 95% credible interval of the mean. For Datasets 1 and 2, this estimated mean falls within a zone of indifference between prospective and retrospective WM, whereas for Dataset 3 the estimated mean falls within a region in which retrospective WM produces superior performance. Higher forgetting rates correspond to more rapid loss of information from WM.

This analysis revealed three notable features of the data. The first is that in all three datasets, every mouse used either a prospective WM code, a retrospective WM code, or both. This result is to be expected given that all animals were trained to criterion on the task, but nevertheless confirms that model M9 provided a good fit to the behaviour of individual animals as well as of group-level behaviour. Second, an overwhelming majority of animals in all three datasets (95% of mice in Dataset 1, 100% of mice in Dataset 2, 83% of mice in Dataset 3) showed evidence of either distal-response biases, side biases, or both. This indicates that these idiosyncratic response biases were a pervasive feature of animal behaviour on the TUNL task.

Third, this analysis revealed that the distribution of prospective and retrospective WM coding was similar across animals in Dataset 1 and Dataset 2, but that animals in Dataset 3 showed a qualitatively different pattern of effects. Within Datasets 1 and 2, a large majority of mice showed effects of prospective WM (95% of animals in Dataset 1, 93% of animals in Dataset 2), whereas the proportion of animals using retrospective WM was significantly smaller in these datasets: 51% of animals in Dataset 1 (*χ*^*2*^(1) = 39.51, *p* < .001, two-sample test for equality of proportions) and 52% of animals in Dataset 2 (*χ*^*2*^(1) = 16.56, *p* < .001). In Dataset 3, by contrast, this difference was significant in the opposite direction: only 31% of animals showed an effect of prospective WM, but 100% showed an effect of retrospective WM (*χ*^*2*^(1) = 35.30, *p* < .001). Simultaneous use of prospective and retrospective WM was relatively common within individual animals in all three datasets (although Dataset 3 once again showed somewhat discrepant results). Within Dataset 1 and 2, relatively few mice used only retrospective WM (5% and 7% respectively), whereas a plurality of mice used either prospective WM only (49% and 47% respectively), or both retrospective and prospective WM (46% and 45% respectively). By contrast, in Dataset 3 the majority of animals used only retrospective WM (69%), none used only prospective WM, and 31% used both retrospective and prospective WM.

The most likely explanation for the discrepant results across datasets is that animals were tested on different trial types in the different datasets: specifically, mice in Dataset 3 were only tested in Separation-1 trials involving the central response location (see Table 1), which is a much smaller subset of trial types than were assessed in Datasets 1 and 2. We therefore sought to ensure that the differing model comparison results between datasets were not a statistical artefact of this difference by re-fitting models to a subset of trials in Datasets 1 and 2 that corresponded to the subset of trials tested in Dataset 3. If the difference between datasets were solely a statistical aberration due to differences in the set of trials tested, then we would expect this subset of trials to show very similar response factor distributions in Datasets 1 and 2 to what we observed in Dataset 3. However, this was not the case (see Supplementary Figure S3), and we therefore conclude that the differences in distributions of response factors across datasets were not driven by methodological differences. Instead, between-datasets differences in the distributions of response factors are likely to reflect the use of different patterns of response factors by mice in Dataset 3 compared with Datasets 1 and 2.

### Prospective WM coding may reflect rational allocation of cognitive resources

The results presented in the previous section raise two important questions: first, given that prospective WM is a suboptimal strategy on the TUNL task, why might so many animals have used prospective memory instead of retrospective WM? Second, why might the distribution of response factors across animals have been so markedly different in Dataset 3 compared with Datasets 1 and 2? These differences might merely reflect unexplained variance within specific animal cohorts or testing facilities; however, another possibility is that the different sets of trial types employed in the different studies differed in the degree to which they gave mice an incentive to use prospective versus retrospective WM.

Here we propose that these two questions can both be answered by considering the balance between prospective and retrospective WM in each dataset in terms of animals’ adaptive responses to the behavioural and cognitive demands of the specific configurations of trial types that were tested. This is consistent with a resource-rational perspective, in which animals’ behaviour is understood as reflecting a trade-off between the competing demands of reward maximisation and physical/cognitive effort minimisation (e.g., Lieder & Griffiths, 2020). We specifically propose that, across all datasets, retrospective WM was more cognitively effortful (i.e., required animals to use more cognitive resources) than prospective WM. In Dataset 1 and 2, mice might therefore have favoured prospective WM as the less effortful of the two coding schemes (despite its lower expected reward). By contrast, the greater relative reward to be obtained^3^ via retrospective WM in Dataset 3 might have given mice in this dataset an incentive to use retrospective WM despite the greater cognitive demands of this response factor.

There are three different lines of evidence in support of this hypothesis: one theoretical, one empirical, and one based on simulations of model performance. From a theoretical perspective, prospective WM is likely to require fewer neurocognitive resources than retrospective WM in the TUNL task because there are fewer possible distinct items that might be coded in prospective WM (two possible prospective response directions vs. five possible retrospective sample locations), therefore requiring less memory allocation. Moreover, forgetting may have been more rapid in retrospective WM because of greater interference between representations of the five different sample locations, compared with the lesser similarity of the two diametrically opposed response directions in prospective WM (Bunting, 2006; Wickelgren, 1965). This would be consistent with Figure 3A, in which the effects of prospective WM take approximately 15 seconds to return to chance-level performance.

This resource-rational explanation makes testable predictions for the data. If it was harder for animals to maintain retrospectively coded items in WM than prospectively coded items, we would also expect that information should be lost more rapidly from retrospective WM than from prospective WM. To test whether this was the case, we formulated an additional set of computational models in which the forgetting rate parameter *β* was allowed to vary between retrospective and prospective WM. The results of this additional set of model comparisons (see Supplementary Table S4) provided evidence that, in line with our resource-rational explanation, information was lost more rapidly from retrospective WM than from prospective WM in Dataset 1 (difference in means of group-level *β* distributions = 0.38; 95% Bayesian HDI: [0.01, 0.86]) and Dataset 2 (mean difference = 3.41; 95% Bayesian HDI: [2.46, 4.53]), though this difference was not credibly different from 0 in Dataset 3 (mean difference = -0.21; 95% Bayesian HDI: [-1.23, 0.41]).

Finally, simulated model performance provides further evidence that animals’ use of prospective WM may have been rational. If information can be held for longer in prospective WM than in retrospective WM, this may mean that animals can acquire more reward using prospective WM at longer retention intervals. This is illustrated via computational simulations of the different WM coding schemes in Figure 5. In particular, the empirical forgetting rates in the best-fitting model in each dataset reveal the resource-rationality of animals’ observed strategy: under the observed WM forgetting rates (i.e., *β* parameters) in Dataset 1 and 2, there was no substantial reward advantage to be gained by using retrospective WM. In Dataset 3, by contrast, for the observed empirical WM forgetting rate, animals could obtain more reward using retrospective WM than using prospective WM. This resource-rational analysis can therefore explain why animals in Dataset 3 used primarily retrospective WM, even as animals in Datasets 1 and 2 used a mixture of both WM types.

The Supplementary Material includes further discussion of the effect of different trial types on the incentives for animals to use prospective vs. retrospective WM, including recommendations on the specific trial types that best isolate retrospective WM in the TUNL task. In short, our simulations suggest that use of trial types at small spatial separations (S0 and S1) promote use of retrospective working memory, whereas trial types at larger spatial separations (S2 and S3) will tend to promote use of prospective working memory. As such, we suggest that probe-phase trials at small spatial separations should be used if experimenters wish to promote use of spatial working memory (as distinct from memory for a prospective behavioural response) in mice completing the TUNL task.

## Discussion

By automating large-scale behavioural data collection, touchscreen tasks represent a paradigm shift in the assessment of translational cognitive phenotypes in non-human animals (Bussey et al., 2012; Oomen et al., 2013). A crucial outstanding question, however, is whether animal behaviour on these tasks has *external validity*: that is, whether tasks measure cognitive processes in animals that are equivalent to cognitive processes in humans, including those that that are aimed to be modelled in humans, including those that are impaired in psychiatric disorders (Pound & Ritskes-Hoitinga, 2018). In this project, we used computational modelling to investigate the cognitive processes that drive behaviour on the touchscreen TUNL task of spatial WM in mice. Our goal was to exploit the rich data produced by the touchscreen paradigm to dissect TUNL behaviour into its cognitive subprocesses, thereby to appraise the extent to which the TUNL task measures an analogue of human spatial WM. Contrary to the common assumption that the TUNL task is solely a measure of spatial WM, we found that behaviour was consistent with a mixture of two distinct WM codes: retrospective WM and prospective WM. In addition, we found that the best-fitting model consistently included two types of response bias, representing individual differences in preferences for responding at particular areas of the testing chamber, independent of the location of the sample stimulus.

Across three separate datasets, our computational modelling results consistently showed that behaviour was best explained by a combination of retrospective WM and prospective WM. That is, the behaviour of mice was consistent with a WM representation that was a mixture of (a) a retrospective representation of the prior spatial location of the sample stimulus, and (b) a prospective representation of the animal’s intended future behavioural strategy (responding at the leftmost vs. the rightmost of the two available response options). This finding stands in contrast to standard TUNL analysis approaches, which implicitly assume that behaviour in this task solely reflects WM for the spatial location of the sample stimulus (i.e., a purely retrospective WM code). In our results, the proportion of behavioural variance that was uniquely accounted for by retrospective WM was never greater than 22% in any of our three datasets, and was as low as 3.1% in Dataset 1. We therefore conclude that in mice, performance on the TUNL task does not solely reflect spatial WM; as such, great care is needed to distinguish behaviours produced by retrospective WM from behaviours produced by other response factors. This is particularly pertinent in translational neuroscience research using this task, where it is critical to ensure that impairments observed in a given genetic, pharmacological or acquired model of a psychiatric or neurological disorder do indeed reflect deficits in retrospective WM, thus capturing clinically-relevant symptoms. Our results suggest the necessity of distinguishing this possibility both from deficits in prospective WM and from changes in the balance between animals’ usage of retrospective vs. prospective WM codes.

A robust behavioural finding, both in the literature and across the datasets that we analysed here, is that animals’ response accuracy reduces with decreasing spatial separation between the sample location and the non-matching response location. This finding has previously been taken as evidence that animals use retrospective WM to complete the task (e.g., Sbisa, Gogos & van den Buuse, 2017; Gogos et al., 2020; Kim et al., 2015). However, our results show that the same qualitative pattern can also be produced by prospective WM, since closer spatial separations also result in greater prospective response ambiguity for an animal using prospective WM (Supplementary Figure S1). Moreover, the datasets that we studied here also showed distinctive behavioural signatures of prospective WM that cannot be explained by retrospective WM, such as significantly below-chance performance on certain S0 trial configurations (Figure 2A). Use of prospective WM may also explain other behavioural phenomena that have been observed in the TUNL literature, such as differences in response accuracy between ‘centre’ and ‘non-centre’ trials (i.e., worse performance for trials in which the sample location is in the centre; Kim et al., 2015). In line with this hypothesis, in Dataset 3 we found that a task design that maximised the number of ‘centre’ trials increased animals’ use of retrospective WM (and decreased their use of prospective WM) relative to Datasets 1 and 2, in which ‘centre’ trials were no more common than any other trial type.

Overall, therefore, our results provide strong evidence that animals performing the TUNL task frequently used prospective WM, rather than solely relying on retrospective WM, as was previously assumed. To explain these findings, we can adopt a resource-rational framework (Lieder & Griffiths, 2020), which considers the cognitive effort costs associated with different behavioural strategies as well as the reward that can be obtained via each strategy. We hypothesise that mice completing the TUNL task used prospective WM because it was less cognitively demanding than use of retrospective WM. As a consequence, we suggest that mice preferred to adopt the less effortful approach of using prospective WM—rather than the more effortful approach of retrospective WM—even though more reward could be obtained via retrospective WM. This explanation can account both for the surprisingly high usage rates of prospective WM in a task that was designed to elicit retrospective WM, as well as for the finding that (in Datasets 1 and 2) information was forgotten more rapidly from retrospective WM than from prospective WM.

To the best of our knowledge, the distinction between prospective and retrospective WM has not previously been considered in the TUNL task. However, this dichotomy has a long history of study in the animal behaviour literature (e.g., Cook et al., 1985; Kametani & Kesner, 1989; Kesner, 1989; Rainer et al., 1999). Consistent with the findings that we report here, rats completing a radial maze task adopted a mixture of prospective and retrospective WM codes depending on the cognitive demands of the task (Kesner, 1989). Human participants also adjust their usage of prospective and retrospective WM depending on task demands (Nallan et al., 1991). However, for the spatial WM tasks of which the TUNL task is putatively an analogue, humans primarily use a retrospective WM code (for review see Zimmer, 2008). As such, in the interests of optimising the external validity of this task, it is important to consider how TUNL task design can be optimised to maximise animals’ use of retrospective WM (see also recent work on a 6-location version of the TUNL task by Dexter et al., 2022).

The TUNL task uses a relatively large set of response option configurations (i.e., unique pairs of sample and non-match location; Figure 1B). For some of these configurations, an animal can respond correctly by using either prospective or retrospective WM; for others, the animal can only consistently respond correctly if it uses retrospective WM, and will perform at or below chance level if it uses prospective WM. As such, it might be possible to promote the use of retrospective WM by increasing the proportion of probe-phase trials that test trial types which can only be solved using retrospective WM. One way of doing this would be to test only S1c trials (i.e., those in which either the sample location or the non-matching location was in the centre): in Dataset 3, which adopted this approach, we found that animals primarily used retrospective WM. More broadly, our simulation analyses (see Supplementary Figure S4) illustrate that testing animals only on S0 trials, or on a combination of S0 and S1 trials, would be expected to have the same effect. By contrast, our results suggest that testing primarily S2 and S3 trials would be expected to increase reliance on prospective WM. Taken together, our results suggest that researchers using the TUNL task may be able to increase the external validity of this task by carefully choosing test-phase trial types that ‘nudge’ animals towards using retrospective WM.

We also found evidence for two distinct types of animal-specific response biases: side biases and distal-response biases. Side biases, which represent animals’ stable and idiosyncratic preferences for responding in a leftward or rightward direction, have been consistently observed in the animal behaviour literature (e.g., Broschard et al., 2021; Kuwabara et al., 2020; Treviño, 2015). As such, it is relatively unsurprising that side biases were also evident in the TUNL task. By contrast, to our knowledge distal-response biases—in which individual animals have stable preferences for responding either closer to the walls or closer to the centre of the testing chamber—have not been previously documented in touchscreen behavioural tasks. We speculate that distal-response biases in the TUNL task may be related to the preference for proximity to vertical surfaces that is observed in many small mammals, including mice, and that is thought to reduce predation risk in naturalistic contexts (e.g., Jensen et al., 2003). In line with this interpretation, we observed a significant group-level preference for distal response options (i.e., those closer to the walls of the chamber) over central response options in two of the three datasets that we analysed. Interestingly, however, in each dataset we observed a small subset of mice (ranging from 7% to 19% of the total sample) that preferred to respond at central response locations. It is possible that this may reflect inter-animal variability in an anxiety-related phenotype, similar to that measured in the elevated plus maze (Walf & Frye, 2007).

From the perspective of studying spatial WM, each of these response biases are important in accounting for TUNL behaviour insofar as they can each be considered a kind of regressor of no interest. These idiosyncratic response biases are likely to be unrelated to the primary cognitive process of interest in the TUNL task since, in most cases, the strength of these response biases was unrelated to overall performance levels on the task. In the absence of a plausible method for abolishing biases from animal behaviour, we suggest that computational modelling can be used to effectively ‘regress out’ these biases from analyses of behaviour.

Several limitations of our computational modelling approach should be noted here. First, the TUNL task is used to assay spatial WM in both mice and rats, but the analyses presented here focus only on behaviour in mice. Notably, the rat TUNL task uses a 3 × 5 grid of possible response locations (Oomen et al., 2013; Sbisa, Gogos & van den Buuse, 2017; Lee et al, 2020), which is more complex than the 1 × 5 grid tested in mice. We consider it plausible that our results may also extend to rat behaviour on the TUNL task; however, this is an empirical question that should be addressed in future research, and more complex models may be required to account for the greater complexity of the spatial grid in the rat version of the task. Second, computational models were fit to aggregate data across animals, and as such do not shed light on the trial-by-trial dynamics of animals’ WM representations. One possibility is that animals strategically used prospective WM for some trial types and retrospective WM for other trial types. An alternative possibility is that, for each animal, behaviour involved a mixture of prospective and retrospective WM in a consistent way across different trial types. Our modelling results cannot discriminate between these two potential explanations; as such, further within-trial data (e.g., *in vivo* neural recordings from hippocampus or prefrontal cortex) is required to adjudicate between these two possibilities. Finally, because mice were trained using a standard training protocol in each of the datasets that we analysed here, our results do not shed light on the extent to which animals’ use of prospective WM might be an artefact produced by the training protocol itself. The training phase of the TUNL task typically begins with the easier high-separation trials and only proceeds to the more difficult low-separation trials after animals reach criterion on the earlier trials. Because retrospective WM is only advantageous over prospective WM for low-separation trials, this training protocol may inadvertently have the effect of increasing the salience of the prospective WM strategy. An open question is whether changes in the training phase of the task, including those recently suggested by Dexter et al., (2022) might alter the balance between prospective and retrospective WM during the subsequent test phase.

In summary, across three distinct datasets our computational modelling results provide evidence that the behaviour of mice on the TUNL task is more multifaceted than has previously been appreciated. We found that retrospective WM, which has previously been assumed to be the dominant factor underlying TUNL performance, only accounts for a portion of the variance in the data. Indeed, in two of the three datasets that we studied, prospective WM was a more significant factor in mouse behaviour than retrospective WM. Our results suggest that this pattern may emerge as the result of a resource-rational trade-off in animals’ behaviour between reward maximisation and cognitive effort minimisation. Moreover, our results suggest a number of factors that might be adjusted in future research using the TUNL task in order to maximise its external validity as a measure of spatial WM. This would be crucial in optimising the utility of the TUNL task as a translational assay of cognition in models of cognitive deficits in neurodevelopmental and psychiatric disorders like schizophrenia, autism, and ADHD.

## Materials and Methods

### Data collection

#### Animal sample

Animal data were derived from three cohorts of mice (Nakamura et al. 2021; Vinnakota et al. in preparation; Sokolenko et al. 2020). All animal experiments were conducted in accordance with the guidelines in the Australian Code of Practice for the Care and Use of Animals for Scientific Purposes (National Health and Medical Research Council of Australia, 8th edition 2013) and the ARRIVE guidelines. Mice from Nakamura et al were approved under La Trobe University Animal Ethics Committee. C57BL/6J mice were obtained from the Animal Resources Centre (Western Australia) and all subsequent husbandry, housing, and touchscreen testing was undertaken at the La Trobe University Animal Research and Teaching Facility (Bundoora, Victoria, Australia). Mice were housed in a reverse 12:12 h light cycle in individually ventilated cages (GM500, Tecniplast, Italy) with *ad libitum* access to food and water until the initiation of food restriction (see below). Cages were monitored daily and changed fortnightly. Mice from Vinnakota et al. were approved under Monash University animal ethics committee (#E/1837/2018/M). Mice were housed in reverse 12:12 h light cycle in open air cages with *ad libitum* access to food and water until initiation of food restriction (see below). C57BL/6J mice were obtained from the Animal Resources Centre (Western Australia) and all subsequent husbandry, housing, and touchscreen testing was undertaken at Monash University, Alfred Health. Mice from Sokolenko et al. were approved under the Florey Neuroscience Institute Animal Ethics Committee (#16-028). C57Bl/6 mice were originally obtained from Jackson Labs, and group-housed in open top cages in the Kenneth Myer Building, Parkville, VIC, Australia, on a reverse-lighting schedule (lights off from 0700-1900). Mice were provided *ad libitum* access to standard chow until 10 weeks of age, when food restriction was initiated (below).

In all datasets, we report analyses from the entire sample of mice without reference to distinctions between experimental groups in the original publications (e.g., maternal immune activation for a subset of mice in Dataset 1). This was because our goal was to assess patterns of behaviour that were observed consistently across multiple datasets of TUNL data from different samples of mice, without reference to distinctions between groups of mice in any specific experiment. Nevertheless, in order to ensure that our results were not driven by impaired performance induced by experimental manipulations, we repeated all analyses among control mice only. The results of these analyses indicated that the same overall behavioural patterns were observed even when we restricted analyses to control mice only (see Supplementary Table S6 and Supplementary Figure S5).

#### Food restriction, habituation and training for touchscreen task

Prior to touchscreen testing, mice were gradually food restricted over a period of approximately 3-5 days until reaching 90% of their free-feeding weight, which was maintained until the end of testing. Strawberry milk (Nippy’s, Australia) liquid food reward was introduced two days before testing to familiarise the mice with the reward. Mice were handled daily for 1 week prior to initiating touchscreen training to habituate to the handler. The TUNL task was run in isolated touchscreen operant chambers for mice (Campden Instruments Ltd, UK) through ABET II software. Chambers were dedicated to either male or female mice only. The apparatus was cleaned with ethanol 80%v/v after each session. The training schedule for mice to learn how to use the apparatus and then to learn the task sufficiently to be tested on the probe trials is provided in Supplementary Table S5.

#### TUNL task

The TUNL task is a touchscreen spatial non-match-to-sample task that was designed as an assay of rodent WM (Bussey et al., 2012; Oomen et al., 2013; Talpos et al., 2010). In each trial of this task, mice must first touch an illuminated ‘sample’ location (one of five horizontally spaced locations on the touchscreen, signalled by increased luminance; see trial schematic in Figure 1A). After a retention interval in which no locations are illuminated, animals complete a choice phase in which two locations are illuminated: one matching the sample location, one non-matching. Animals receive a reward if they touch the non-matching location. In Datasets 2 and 3 (but not Dataset 1), incorrect responses were followed by ‘correction trials’, in which the same two choice locations remained illuminated until animals correctly selected the non-matching location.

This task is considered a test of spatial WM because optimal task performance requires the animal to maintain the spatial location of the sample in WM during the retention interval, so that the non-matching location can be selected at the choice phase. Trial difficulty in the TUNL task is manipulated either by increasing the duration of the retention interval (thereby increasing the delay over which the sample location must be held in WM), or by reducing the spatial separation between the sample location and the non-matching location (thereby increasing the similarity between the two possible choice options). Different separation conditions in the TUNL task are typically described based on the number of unlit locations that separate the two response options in the choice phase (e.g., the example trial in Figure 1A is referred to as a Separation-3 or S3 trial because three unlit squares separate the two response options in the choice phase). In the present study, we solely analysed data from the ‘probe’ phase of the task, after animals had learned the task to criterion. All mice were trained to criterion on the task using a standard habituation and conditioning protocol (Kim et al., 2015).

### Data analysis

#### Logistic regression analyses

The effects of separation, delay, and trial type on behaviour on the TUNL task were assessed using mixed-effects logistic regression analyses, with choice accuracy as the dependent variable (choice of non-match location coded as 1, choice of sample location coded as 0). Analyses were carried out using the *lme4* package for R (Bates et al., 2015). All regression models included per-animal random intercepts, as well as per-animal slopes for all main effects and interactions that were entirely within-animal (Barr et al., 2013). Non-converging models were simplified by removing higher-order random slopes until convergence was achieved. Coefficient *p*-values were calculated using the Wald t-to-z test (Meteyard & Davies, 2020).

#### Calculation of side bias metric

We computed a model-agnostic measure of the strength of each animal’s preference for left/right response options, taking into account the different choice options available within the different datasets. To do this, we first calculated the *choice disparity* for each of the five possible response locations, which is a normalised measure of how often each animal responded at each response location (cf. Broschard et al., 2021):

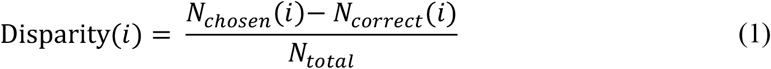

Here, *i ∈* [1, 2, 3, 4, 5] denotes the response location (1 = far left, 3 = centre, 5 = far right), *N*_*chosen*_(*i*) denotes the number of times that the animal responded at location *i* and *N*_*correct*_(*i*) denotes the number of times that location *i* was the correct response location. The denominator *N*_*total*_ denotes the total number of trials completed by the animal, allowing us to compare the choice disparity index across animals who completed different total numbers of trials. This metric can be interpreted as an index of whether an animal chose a particular response location more often than expected given the trial types it completed (positive disparity), or less often than expected (negative disparity).

For each animal, we then computed the side bias index simply by calculated the difference between the choice disparity for rightward response options and the choice disparity for leftward response options. This provides a normalized index of whether an animal preferred rightward response options (positive side bias metric) or leftward response options (negative metric):

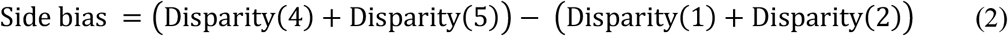

Within Figure 3, we computed the animal-by-animal significance of the side bias metric using a non-parametric empirical permutation test. For each animal, we tested significance by randomly shuffling the actual *N*_*chosen*_ vector 1000 times to estimate an empirical null distribution of the side bias metric. An animal’s side bias was taken to be significantly different from chance if the actual metric as computed by Equation 2 fell outside the 95% confidence interval of this metric as estimated by the empirical permutation test.

### Calculation of distal-response bias metric

The computation of the distal-response bias metric was also based on differences in choice disparities as defined in Equation 1. In this case, however, we were solely concerned with whether each animal responded at the central response location more or less often than would be expected by chance:

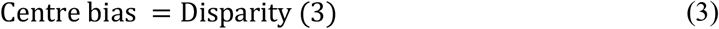

This computes an index of whether animals preferred the central response option or more distal response options (positive/negative values respectively). As with the side bias index, significance was estimated on a per-animal basis using a non-parametric empirical permutation test.

### Computational models

All models assumed that an animal’s choice on a given trial was driven by a competition between two response strengths (cf. Kruschke, 1992; Nosofsky, 1986): one for the matching location and one for the non-matching location (denoted *R*_*m*_ and *R*_*nm*_ respectively). In all models, the probability of making a (correct) non-match response was a logistic function of the difference between these competing response strengths (also known as a softmax function or Luce choice rule) on a given trial *t*:

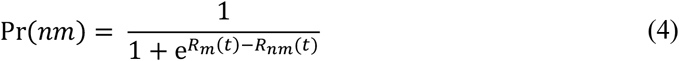

Different models made contrasting assumptions about how these response strengths were calculated. Specifically, different models comprised different additive combinations of four different response factors: retrospective WM, prospective WM, side biases, and distal-responses biases. In Model M9, for instance, all four response factors influenced choice, and response strengths were calculated as per Equation 5:

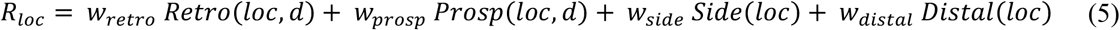

Where *loc* stands for either the sample location *m* or the non-matching location *nm* (*m, nm ∈* [1, 2, 3, 4, 5]), and *d* denotes the duration of the retention interval. Each of the *w* parameters in Equations 5 and 6 is an animal- and component-specific weighting parameter that controls the strength of each of the response factors (as specified by the functions *Retro, Prosp, Side*, and *Distal*; see below). The simpler models M1-M8 consisted of more restricted combinations of the different combinations, as specified in Table 2. For instance, response strengths in the retrospective-memory-only M1 were computed as per Equations 7 and 8:

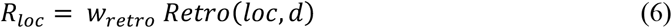

In all models, the key distinction between the working-memory factors (*Retro* and *Prosp* functions) and the response-bias factors (*Side* and *Distal*) was that the response strength produced by working-memory factors was assumed to decrease over time as information was forgotten from WM (i.e., these functions depend both on the locations *m* and *nm* and the delay *d*), whereas the response strength produced by response biases was assumed to remain constant over time (and therefore depend only on *m* and *nm*). In all models reported in the main text, we modelled the rate of this information loss using a power-law forgetting function (Donkin & Nosofsky, 2012; Wickelgren, 1974; Wixted & Carpenter, 2007):

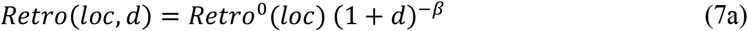

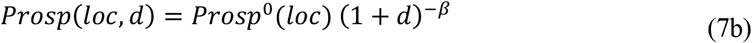

Here, the remaining response strength produced by WM after *t* seconds depends both on the initial response strength functions *Retro*^0^ and *Prosp*^0^ (i.e., the response strength immediately after the presentation of the sample location as per retrospective and prospective WM respectively; see below) and the retention duration *d*. The free parameter *β* controls the rate at which information is forgotten from WM^4^.

We also tested several alternative forgetting functions (exponential and logistic/sigmoidal functions; Equations 8 and 9 respectively). However, we found the power-law function provided the best overall fit to data, and the rank order of different models’ goodness of fit was largely unchanged across the different forgetting functions (see Supplementary Tables S1 and S2). We therefore solely report conclusions from power-law computational models in the main text.

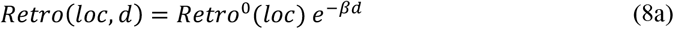

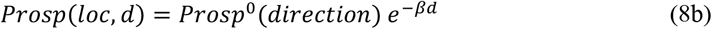

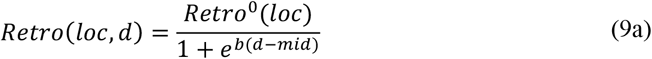

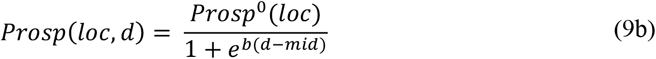

*Retrospective WM*. The retrospective WM equation instantiated the hypothesis that animals made choices in the TUNL task by holding the location of the original sample stimulus in WM during the retention interval. At the choice phase, each possible response location was then assumed to accrue response strength in proportion to its spatial distance to the sample location (i.e., greater response strength for locations at a greater distance from the sample location). We modelled this accrual of response strength using an exponential similarity kernel (Shepard, 1957, 1987):

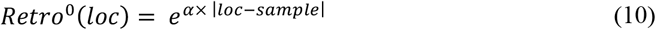

Where *α* is a free parameter corresponding to the kernel width of the dissimilarity kernel. This parameter can be interpreted as the degree of cognitive dissimilarity between different response locations (i.e., cognitive capacity for pattern separation). When *α* is large, different response locations are sharply distinguished from one another; as alpha approaches 0, the animal treats all possible response locations as equivalent. To avoid parameter non-identifiability, the *α* parameter was constrained to be positive in all datasets. *α* was fixed to a value of 1 in Dataset 3, because the paucity of different separation conditions in this dataset (see Table 1) meant that the *α* parameter was not uniquely identified.

### Prospective WM

The prospective WM equation instantiated the hypothesis that animals made choices in the TUNL task by holding an intended response direction in WM during the retention interval. That is, upon seeing the sample location, animals were assumed to form a prospective behavioural intention to select either the leftmost or the rightmost response location at the choice phase (according to whichever direction was most associated with reward in training for each sample location; see Table 1 and Table S1). Specifically, this model component assumed that the response strength for the different locations at the choice phase was proportional to the probability that that response direction would be correct given the sample location:

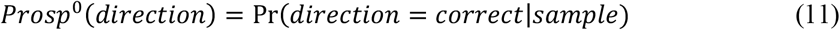

These probabilities were assumed to be learned by animals throughout the conditioning phases of the task.

*Distal-response bias*. This response factor assumed that, independent of all other task factors, individual animals idiosyncratically preferred to respond either closer to the walls of the testing chamber (both left and right walls), or to respond closer to the centre of the testing chamber:

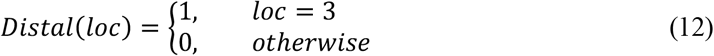

Where *loc* = 3 denotes that the response location was in the centre of the five possible locations. Accordingly, a positive value of the weighting parameter *w*_*distal*_ denotes a preference for central stimuli, a negative value denotes a preference for distal stimuli, and a value of zero denotes no overall preference for central vs. distal response options.

### Side bias

This response factor assumed that, independent of all other task factors, individual animals idiosyncratically preferred to choose either the leftward or the rightward of the two response options on a given trial:

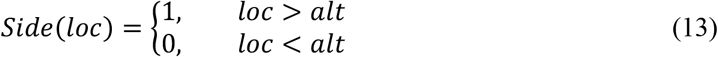

Where *loc* > *alt* indicates that the response location was to the right of the alternative location, and *loc < alt* indicates that it was to the right. Accordingly, a positive value of the weighting parameter *w*_*side*_ indicates a preference for rightward stimuli, a negative value indicates a preference for leftward stimuli, and a value of zero indicates no bias in either direction.

### Model fitting and comparison

All models were fit within a hierarchical Bayesian framework using the probabilistic programming language Stan (Carpenter et al., 2017) and the *cmdstanr* package for R. For each model, we took 3000 samples from the joint posterior per chain across a total of four chains. The first 1750 samples per chain were discarded to prevent dependence on initial values, resulting in a total of 5000 post-warmup posterior samples being retained for analysis across the four chains. Models were compared using the Watanabe-Akaike Information Criterion (WAIC; Watanabe, 2013) as estimated within the *loo* package in R (Vehtari et al., 2017). This criterion is an approximation of the leave-one-out posterior predictive density of the data given the model, and assesses goodness of fit while penalising models with excessive complexity. Statistical ties between different models (i.e., differences in WAIC of less than one standard error of the difference of WAIC) were broken according to statistical parsimony, defined as number of parameters per animal (Table 2). All chains converged in all models as indicated by an 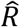 value less than 1.1, and there were no divergent transitions in any model. All models used partial pooling, with animal-level parameters assumed to be drawn from a group-level distribution with a mean and standard deviation estimated from the data. Point estimates of animal-level parameters were calculated as the median of the posterior distribution per parameter per animal.

### Variance partitioning analysis

In order to compute the amount of variance explained by each response factor in the best-fitting model M9, we conducted a variance partitioning analysis. In this analysis, we calculated the change in the overall group-level model *R*^*2*^ when different response factors were removed one at a time from model M9. To compute the proportion of variance uniquely explained by retrospective WM, for instance, we computed the difference in *R*^*2*^ between model M9 and model M5; for prospective WM, the *R*^*2*^ difference between M9 and M6, and so on. *R*^*2*^ values in this analysis were computed for each trial type and delay duration at the group-mean level.

### Model simulations

We conducted two sets of simulation analyses to dissect the behavioural signatures of retrospective versus prospective WM in the TUNL task.

First, we conducted several simple simulations to calculate the expected proportion correct for different configurations of sample location and non-match location under retrospective WM versus prospective WM (Figure 1B and Supplementary Figure S1). In these simulations, we assumed that each of the 20 trial types in Figure 1B was presented equally often, and that animals deterministically selected the response locations indicated by either a retrospective or prospective WM code (or selected at random between the different indicated locations in cases where more than one response location was equally indicated). For retrospective WM, simulations assumed that animals were perfectly able to select the non-matching location. For prospective WM, simulations assumed that animals selected a response direction (leftward versus rightward) on the basis of the sample location alone, and then deterministically responded at the location that was further in the selected response direction in the choice phase. The response direction was assumed to be learned during the conditioning phase of the task by marginalising across the different subsequent non-match locations that followed each possible sample location. For instance, when the sample location was second from the left (second column in Figure 1B), then 75% of the time the correct response location would be the rightward of the two options in the choice phase. The simulations reported in Supplementary Figure S1 therefore assume that mice deterministically responded at the rightward response location after a sample location that was second from the left (and were therefore correct on 75% of trials for this sample location).

The second set of simulations relaxed the unrealistic assumption that animals were able to perfectly execute either retrospective or prospective WM. Here, we estimated the performance of both retrospective and prospective WM under the assumption that the response strength according to WM (of either kind) became weaker over time, consistent with the behavioural data presented in Figure 2. For each dataset, we specifically estimated the performance of each WM code using point estimates of the forgetting rate parameter *β* for the best-fitting model for that dataset (for Dataset 1: model M9 with two distinct *β* parameters; for Datasets 2 and 3: model M9 with a single shared *β* parameter; see Supplementary Table S4). Since these simulations were probabilistic rather than deterministic, we repeated simulations 10,000 times and computed the proportion of datasets in which an animal using retrospective WM alone would obtain more reward than an animal using prospective WM alone. For each dataset, we simulated performance according to the actual configuration of trial types that were presented in that dataset (Figure 5). To inform the design of future studies, we also simulated performance under other trial configurations that experimenters might wish to test (Supplementary Figure S4).

## Footnotes

1 Although prospective WM can produce above-chance TUNL performance overall, this is not true for all TUNL trial types considered separately. For instance, prospective WM cannot be used to respond accurately for a central sample stimulus on the TUNL, because in that case the correct response option at the test phase is equally likely to be the leftmost or the rightmost option (see ‘Sample in centre’ column of Figure 1B). Further simulations of TUNL performance under different memory coding strategies are presented in Supplementary Figure S1

2 In the interests of succinctness, we here refer to ‘distal-response biases’ even though some mice preferred central response locations. In the nomenclature that we adopt here, such a preference would be represented as a *negative* distal-response bias (i.e., a preference *away from* distal response locations)

3 Dataset 3 tested a different subset of trial types to Datasets 1 and 2 (see Table 1 for details). Specifically, Dataset 3 tested solely ‘S1c’ trials, in which prospective working memory cannot produce above-chance performance (see Figure S1). The advantage for retrospective working memory is therefore significantly greater in Dataset 3 than in Datasets 1 and 2.

4 In the additional models reported in Supplementary Table S4, different *β* parameters were fit for Equation 7a and Equation 7b for each animal. In all other models, a single *β* per animal was fit for both 7a and 7b

