## Supplementary Information for "Mouse behaviour on the trial-unique non-matching-to-location (TUNL) touchscreen task reflects a mixture of distinct working memory codes and response biases"

for

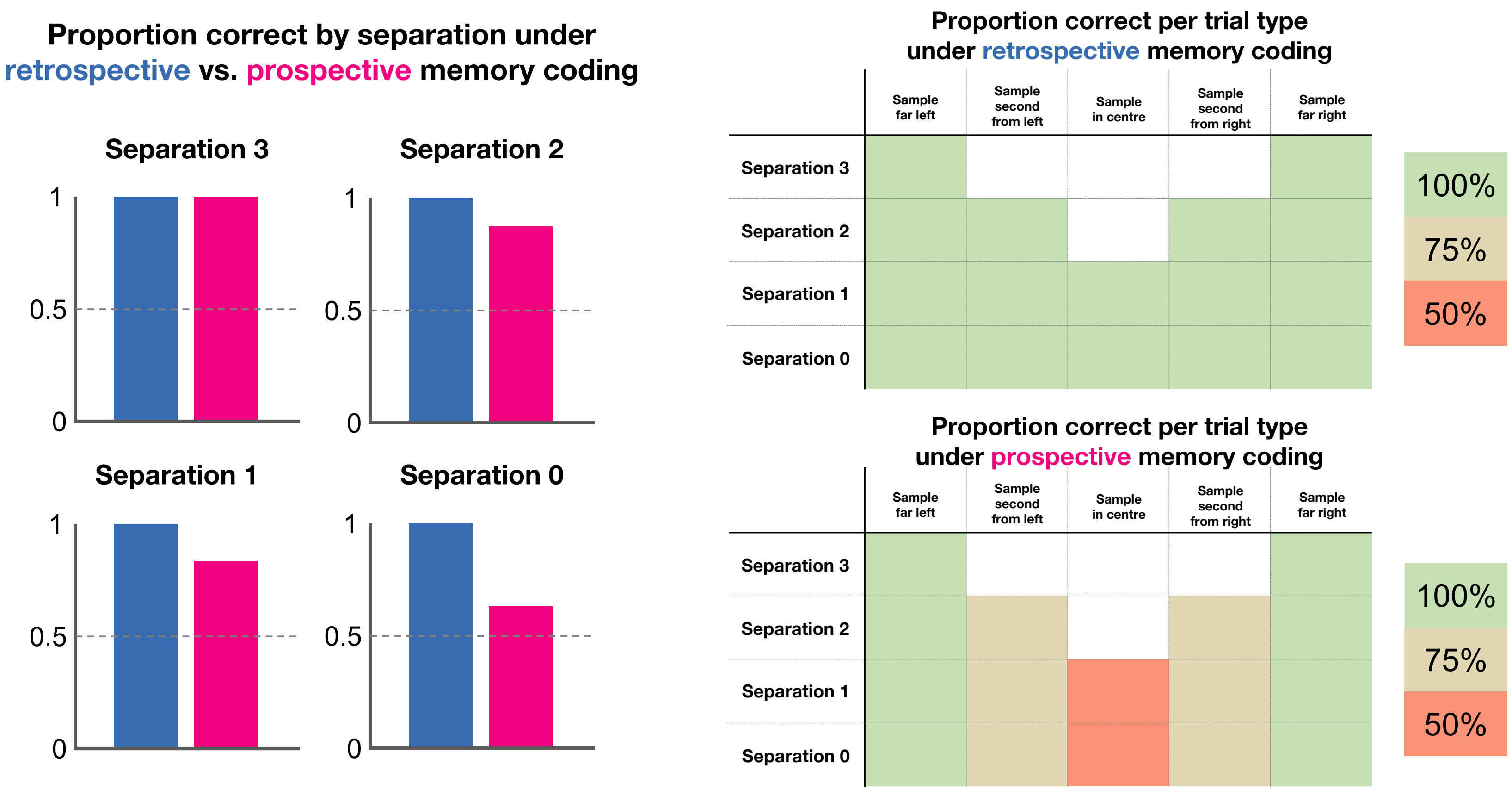


*Supplementary Figure S1.* Simulated proportion of correct responses under retrospective memory coding (remembering the location of the sample stimulus) vs. prospective memory coding (remembering the direction in which to respond after the retention interval). Left: expected proportion of correct responses is 100% under retrospective memory coding in all separation conditions, and is lower under prospective memory coding (though still above-chance in all separation conditions). Right: Expected proportion of correct responses varies as a function of sample location for prospective memory coding, but not for retrospective memory coding. Simulations assume that all trial configurations (laid out as per Figure 1B) are presented equally often, and that both retrospective memory coding and prospective memory coding are errorless.

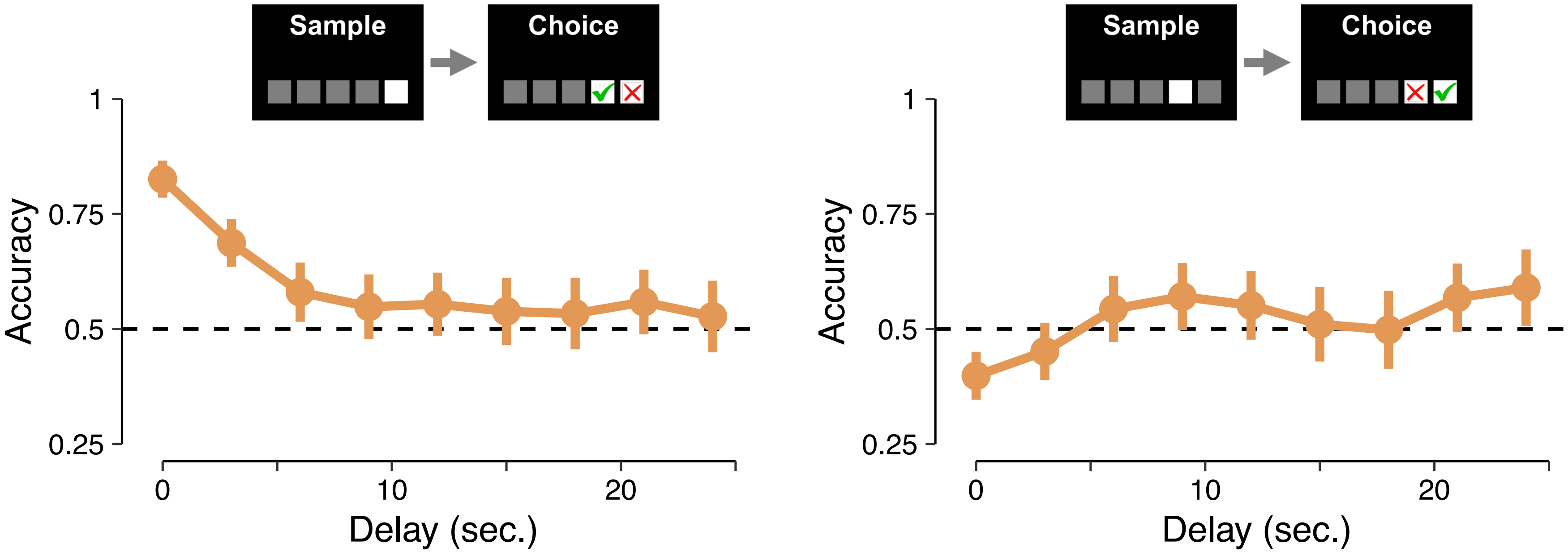


*Supplementary Figure S2.* Demonstration of direction-dependent effects in two putatively similar S0 choice configurations across multiple delays from Dataset 1. In the left subplot, the sample location [red cross] is at the far right and the non-match location [green tick] is second from the right; when the trial is presented in this configuration, mice display above-chance performance at delay 0 (mixed-effects logistic regression; intercept = 1.07, $p < .001$) that deteriorates with increasing delay duration ($\beta = -0.06$, $\chi^2(1) = 32.33$, $p < .001$). In the right subplot, when the sample location is second from the right and the non-match location is at the far right, mice show below-chance performance at delay 0 (intercept = -0.28, $p = .01$) and improving performance with increasing delays ($\beta = 0.04$, $\chi^2(1) = 10.95$, $p < .001$). Error bars represent the 95% confidence interval of the mean.

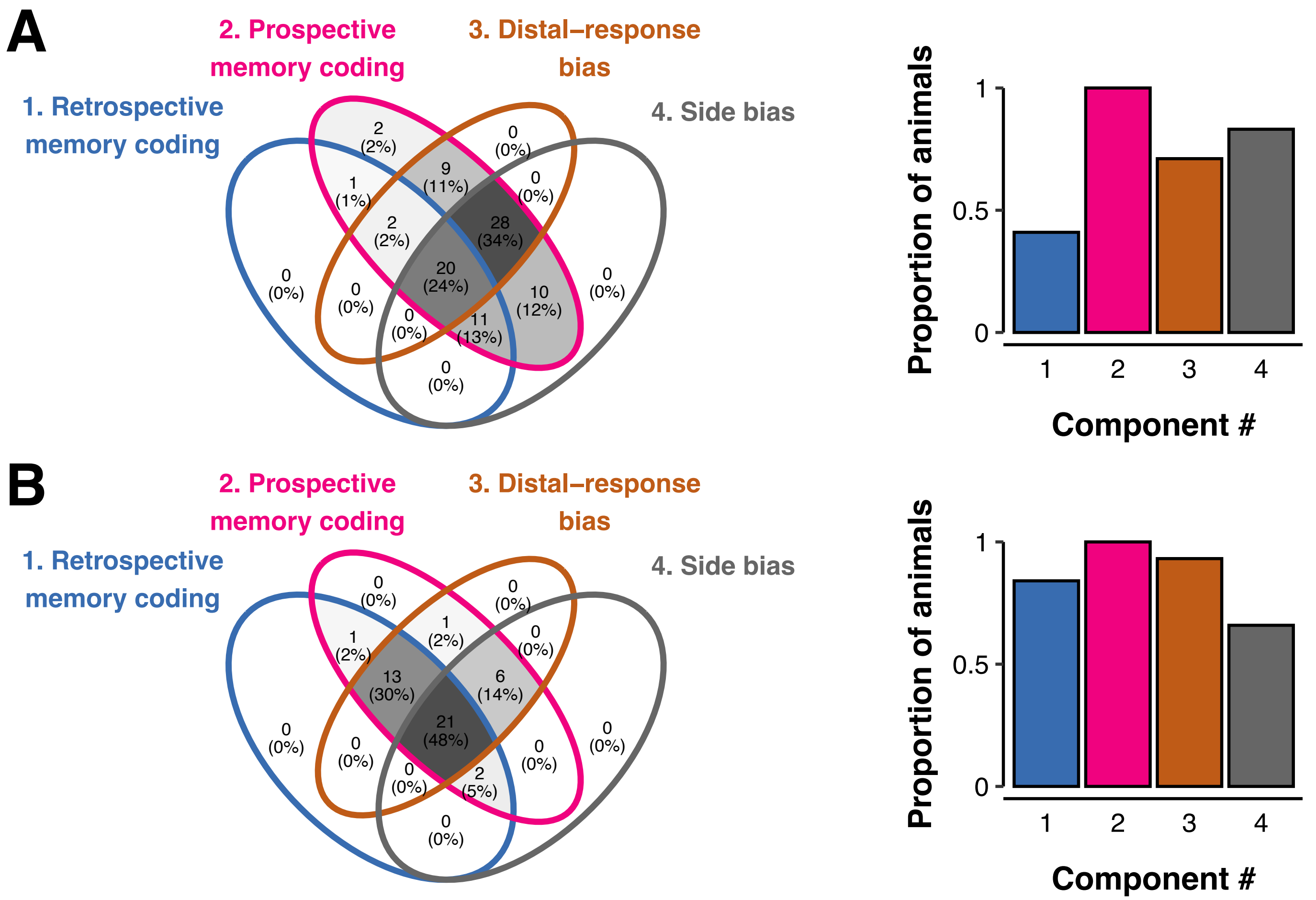


*Supplementary Figure S3.* Breakdown of significant model components across mice in a subset of trials from Dataset 1 (**A**) and Dataset 2 (**B**) designed to match the trial types tested in Dataset 3. Venn diagrams present the breakdown of the numbers of mice who fit into each possible configuration of significant model components (presented as raw numbers and percentage of dataset). Histograms depict the proportion of mice in each dataset who displayed a significant effect of each model component overall (i.e., the sum of the percentages within each coloured circle in the Venn diagrams).

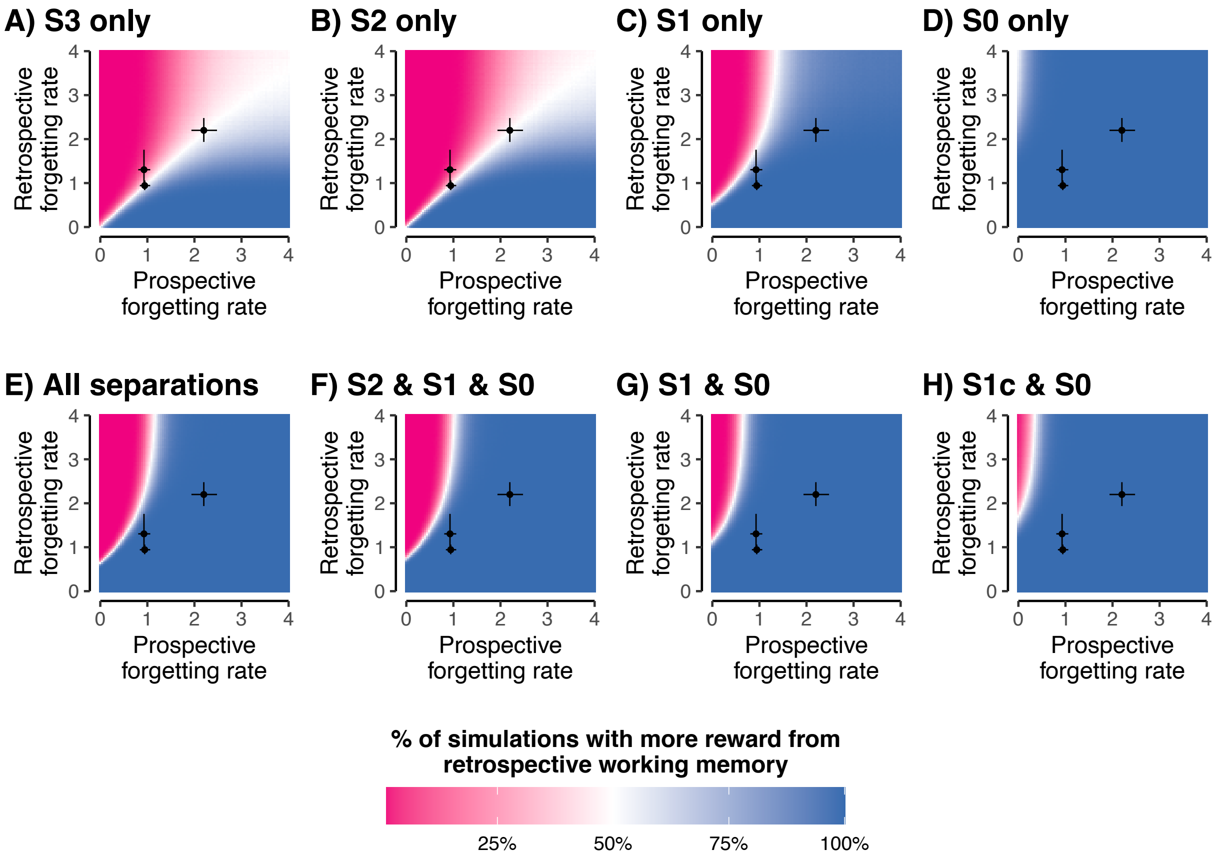


*Supplementary Figure S4.* Simulated relative performance of prospective and retrospective working memory across different configurations of trial types. Blue regions of each heatmap denote parameter regimes under which more reward can be acquired using retrospective working memory, whereas pink regions denote parameter regimes in which prospective working memory is more advantageous. White regions denote parameter regimes in which retrospective and prospective working memory produce equivalent performance on average. Points and error bars denote the mean estimated forgetting rate for each working memory types +/- the 95% credible interval of the mean across each of the three datasets. In general, retrospective working memory is favoured at smaller separations and prospective working memory is relatively more favoured at larger separations. Simulations assume 48 trials per trial type per animal at each of 4 delay durations (0, 2, 4, 6 seconds).

*Supplementary Table S1.* Overview of fits of computational models with an exponential forgetting function to Dataset 1

| Model number | *N* parameters per mouse | Forgetting function | Retrospective working memory | Prospective working memory | Side biases | Distal-response biases | WAIC | ΔWAIC (SE) |
|---|---|---|---|---|---|---|---|---|
| E1 | 3 | Exponential | ✓ | - | - | - | 71,402.8 | 15,866.6 (362.5) |
| E2 | 2 | Exponential | - | ✓ | - | - | 71,333.7 | 15,842.7 (361.0) |
| M3 | 1 | N/A | - | - | ✓ | - | 66,937.5 | 11,308.9 (371.5) |
| M4 | 1 | N/A | - | - | - | ✓ | 76,250.8 | 20,622.3 (422.1) |
| E5 | 4 | Exponential | - | ✓ | ✓ | ✓ | 55,951.0 | 322.5 (61.2) |
| E6 | 5 | Exponential | ✓ | - | ✓ | ✓ | 56,997.2 | 1,368.7 (115.4) |
| E7 | 5 | Exponential | ✓ | ✓ | - | ✓ | 68,122.4 | 12,493.9 (334.3) |
| E8 | 5 | Exponential | ✓ | ✓ | ✓ | - | 58,012.7 | 2,384.2 (152.5) |
| E9 | 6 | Exponential | ✓ | ✓ | ✓ | ✓ | **55,628.5** | **-** |

Note: WAIC values are presented on a deviance scale (lower values indicate better model fit).

*Supplementary Table S2*. Overview of fits of computational models with a sigmoidal forgetting function to Dataset 1

| **Model number** | ***N* parameters per mouse** | **Forgetting function** | **Retrospective working memory** | **Prospective working memory** | **Side biases** | **Distal-response biases** | **WAIC** | **ΔWAIC (SE)** |
|---|---|---|---|---|---|---|---|---|
| S1 | 4 | Sigmoidal | ✓ | - | - | - | 71,495.2 | 15,878.6 (363.2) |
| S2 | 3 | Sigmoidal | - | ✓ | - | - | 71,471.2 | 15,854.7 (362.0) |
| M3 | 1 | N/A | - | - | ✓ | - | 66,937.5 | 11,320.9 (371.6) |
| M4 | 1 | N/A | - | - | - | ✓ | 76,250.8 | 20,634.3 (421.6) |
| S5 | 5 | Sigmoidal | - | ✓ | ✓ | ✓ | 55,989.2 | 372.6 (66.8) |
| S6 | 6 | Sigmoidal | ✓ | - | ✓ | ✓ | 56,971.4 | 1,354.9 (116.0) |
| S7 | 6 | Sigmoidal | ✓ | ✓ | - | ✓ | 68,109.6 | 12,493.1 (334.2) |
| S8 | 6 | Sigmoidal | ✓ | ✓ | ✓ | - | 58,074.6 | 2,458.1 (153.1) |
| S9 | 7 | Sigmoidal | ✓ | ✓ | ✓ | ✓ | **55,616.5** | **-** |

Note: WAIC values are presented on a deviance scale (lower values indicate better model fit).

*Supplementary Table S3*. Summary of model parameters for best-fitting model (M9)

| **Parameter name** | **Modelled process** | **Parameter description** | **Dataset 1** | | **Dataset 2** | | **Dataset 3** | |
|---|---|---|---|---|---|---|---|---|
| | | | **Parameter group mean** | **95% HDI for group mean** | **Parameter group mean** | **95% HDI for group mean** | **Parameter group mean** | **95% HDI for group mean** |
| $w_{retro}$ | Retrospective working memory | Decision weight for output of retrospective working memory | 8.43 | [3.78, 17.33] | 14.48 | [13.67, 38.60] | 0.38 | [0.31, 0.45] |
| $\alpha$ | Retrospective working memory | Spatial similarity function within retrospective working memory | 0.03 | [0.01, 0.05] | 0.06 | [0.03, 0.11] | fixed to 1 | - |
| $w_{prosp}$ | Prospective working memory | Decision weight for output of prospective working memory | 1.71 | [1.51, 1.92] | 8.46 | [6.51, 10.88] | 0.91 | [0.46, 1.37] |
| $\beta$ | Prospective and retrospective working memory | Forgetting rate within working memory | 0.99 | [0.91, 1.08] | 2.20 | [1.94, 2.48] | 0.94 | [0.84, 1.06] |
| $w_{side}$ | Side bias | Decision weight for side biases (negative: prefer left; positive: prefer right) | 0.11 | [-0.04, 0.28] | 0.09 | [-0.01, 0.19] | 0.07 | [-0.04, 0.18] |
| $w_{distal}$ | Distal-response bias | Decision weight for distal-response biases (negative: prefer distal; positive: prefer central) | 0.53 | [0.44, 0.65] | 0.56 | [0.44, 0.69] | 0.19 | [0.08, 0.31] |

*Supplementary Table S4.* Comparison of models with one versus two forgetting rates

| Dataset | WAIC (with one forgetting rate) | WAIC (with two forgetting rates) |
|---|---|---|
| 1 | 55,816.5 | **55,476.4** |
| 2 | **7,355.7** | 7,409.5 |
| 3 | **3,365.8** | 3,372.6 |

Note: WAIC values are presented on a deviance scale (lower values indicate better model fit).

*Supplementary Table S5.* Training protocol prior to data collection

| Schedule | Description of training | Criteria |
|---|---|---|
| Habit 1 (20 min) | Habituation to the chamber for 20 minutes | 2 days |
| Initial touch (30 trials out of 60 min) | Associate stimulus with reward delivery | If 30 trials completed in 60 min, then move to next stage. |
| Must touch (30 trials out of 60 min) | Nose-poke stimulus to receive a reward | If 30 trials completed in 60 min, then move to next stage. |
| Must Initiate (30 trials out of 45 min) | Initiate the next trial by breaking infrared (IR) beam near reward collection tray | If 30 trials completed in 45 min, then move to next stage. |
| Punish Incorrect (48 trials out of 30 min) | Incorrect touch (blank windows) leads to the house light switching on for 5s and the trial being repeated until the mouse makes a correct response. | If overall accuracy over 2 consecutive days reaches > 80% correct then move to next stage. |
| **Separation training Stage 1 (36 trials out of 45 min)** | Sample stimuli presented in non-centre locations, starting with maximal separation level (separation 3) and then decreasing separation until separation 1. | If overall accuracy over 2 consecutive days reaches >70% correct, then move to the next separation level. |
| Exp1Stage1 Separation 3 (S3)<br>Exp1Stage1 S2<br>Exp1Stage1 S1 | | |
| **Separation training Stage 2 (48 trials out of 60 min)** | Sample stimuli presented in centre locations, S1 and S0. | Daily training till group performance stabilises (>80%). |
| Exp1Stage2 S1 (1 separation)<br>Exp1Stage2 S0 (0 separation) | | |
| **Probe trials** | | |

*Supplementary Table S6.* Overview of computational model fits to control animals only in each dataset

| **Model number** | ***N* model parameters per mouse** | **Forgetting function** | **Retrospective working memory** | **Prospective working memory** | **Side biases** | **Distal-response biases** | **Dataset 1** | | **Dataset 2** | | **Dataset 3** | |
|---|---|---|---|---|---|---|---|---|---|---|---|---|
| | | | | | | | **WAIC** | **ΔWAIC (SE)** | **WAIC** | **ΔWAIC (SE)** | **WAIC** | **ΔWAIC (SE)** |
| M1 | 3 * | Power-law | ✓ | - | - | - | 34,850.0 | 6,244.3 (229.8) | 4,322.4 | 414.5 (61.3) | 2087.0 | 254.4 (58.1) |
| M2 | 2 | Power-law | - | ✓ | - | - | 35,029.1 | 6,423.3 (233.0) | 4,569.0 | 661.2 (78.2) | 3998.1 | 2165.5 (284.3) |
| M3 | 1 | N/A | - | - | ✓ | - | 33,837.6 | 5,231.8 (247.9) | 7,763.1 | 3,855.3 (224.8) | 6052.6 | 4219.9 (444.7) |
| M4 | 1 | N/A | - | - | - | ✓ | 37,104.2 | 8,498.4 (278.3) | 7,859.8 | 3,951.9 (221.6) | 6094.8 | 4262.2 (441.3) |
| M5 | 4 | Power-law | - | ✓ | ✓ | ✓ | 28,792.0 | 186.3 (60.7) | **3,907.9** | **-** | 2099.9 | 267.2 (63.5) |
| M6 | 5 * | Power-law | ✓ | - | ✓ | ✓ | 29,129.8 | 524.1 (77.5) | 4,034.9 | 127.0 (36.3) | 1841.0 | 8.3 (13.2) |
| M7 | 5 * | Power-law | ✓ | ✓ | - | ✓ | 33,366.0 | 4,760.3 (207.7) | 4,184.2 | 276.3 (48.2) | 2025.7 | 193.1 (51.9) |
| M8 | 5 * | Power-law | ✓ | ✓ | ✓ | - | 29,802.6 | 1,196.9 (100.9) | 4,026.0 | 118.1 (33.7) | 1865.5 | 32.9 (15.9) |
| M9 | 6 * | Power-law | ✓ | ✓ | ✓ | ✓ | **28,605.7** | **-** | 3,909.3 | 1.4 (6.2) | **1832.6** | **-** |

Note: WAIC values are presented on a deviance scale (lower values indicate better model fit).

* One fewer parameter per mouse in Dataset 3 because a smaller number of separations in this dataset rendered the $\alpha$ parameter non-identifiable (see Method)

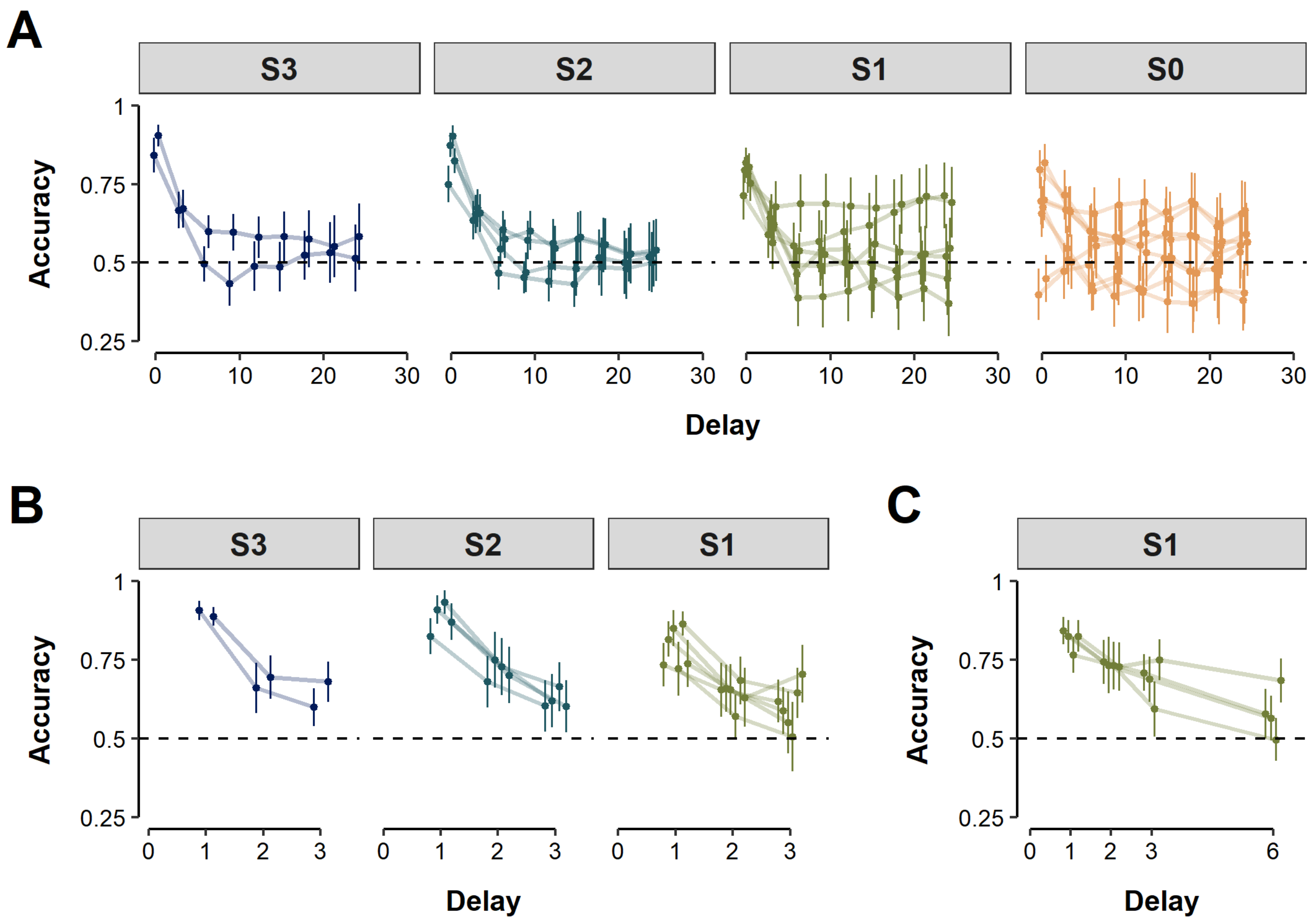


*Supplementary Figures S5.* Mouse behavioural data as a function of separation and delay across wild-type **control mice only** in three independent datasets (**A**: Nakamura et al., 2021; **B**: Vinnakota et al., in prep.; **C**: Sokolenko et al., 2020). Each subplot depicts mean proportion of correct responses (y-axis) across animals as a function of delay between memory sample and response screen (x-axis) and separation condition (plot facets). Each individual point within a delay/separation condition presents data from a unique configuration of sample location and non-match location (see Figure 1B). Error bars represent the 95% confidence interval of the mean.
